# Sequence-dependent mechanochemical coupling of helicase translocation and unwinding at single-nucleotide resolution

**DOI:** 10.1101/2022.02.10.479955

**Authors:** Andrew H. Laszlo, Jonathan M. Craig, Momčilo Gavrilov, Ramreddy Tippana, Ian C. Nova, Jesse R. Huang, Hwanhee C. Kim, Sarah J. Abell, Mallory deCampos-Stairiker, Jonathan W. Mount, Jasmine L. Bowman, Katherine S. Baker, Hugh Higinbotham, Dmitriy Bobrovnikov, Taekjip Ha, Jens H. Gundlach

**Affiliations:** Department of Physics, University of Washington, Seattle, WA, 98195, USA; Department of Physics, Johns Hopkins University, Baltimore, MD, 21218,USA

## Abstract

We used single-molecule nanopore tweezers (SPRNT) to resolve the millisecond single-nucleotide steps of Superfamily 1 helicase PcrA as it translocates on, or unwinds, several kb-long DNA molecules. We recorded over 2 million enzyme steps under various assisting and opposing forces in diverse ATP and ADP conditions to comprehensively explore the mechanochemistry of PcrA motion. Forces applied in SPRNT mimic forces and physical barriers PcrA experiences *in vivo*, such as when the helicase encounters bound proteins or duplex DNA; we show how PcrA’s kinetics change with such stimuli. SPRNT allows for direct association of the underlying DNA sequence with observed enzyme kinetics. Our data reveal that the underlying DNA sequence passing through the helicase strongly influences the kinetics during translocation and unwinding. Surprisingly, unwinding kinetics are not solely dominated by the base-pairs being unwound. Instead, the sequence of the single stranded DNA on which the PcrA walks determines much of the kinetics of unwinding.

## Introduction

Helicases are involved in nearly every aspect of nucleic-acid metabolism throughout the tree of life. A full picture of their mechanochemical action is currently missing because of the difficulty of observing the small fast steps taken by helicases. Helicases use repeated cycles of ATP hydrolysis to generate directed inchworm-like motion along a single-stranded (ss) nucleic acid and thereby unwind nucleic acid duplexes and remove bound proteins(1–4), performing key roles during replication, transcription, and translation of nucleic acids. Biophysical approaches(5) such as X-ray crystallography(6), single-molecule FRET(7), optical tweezers(8–12), and magnetic tweezers(13) have shed light on helicase mechanisms, yet understanding of the mechanochemical choreography of translocation and unwinding is incomplete in part because helicases take small steps (<0.6 nm) at high speed (~1 step per ms), which is beyond the resolution of most single-molecule techniques.

Single-molecule picometer resolution nanopore tweezers (SPRNT, Fig. 1) is a technique for measuring the activities of nucleic acid processing enzymes at high spatiotemporal resolution(14, 15). SPRNT records the motion of ss nucleic acid as a motor enzyme feeds it through a *Mycobacterium smegmatis* porin A (MspA) nanopore (Fig. 1A, Materials and Methods). A voltage applied over the nanopore causes an ion current through the pore and applies an electrophoretic force to the DNA. As the motor enzyme draws DNA through the pore, different bases within the pore block the pore to differing degrees and thereby characteristically affect the ion current. The procession of ion current values is used to determine the nucleic acid sequence which serves as the basis for nanopore sequencing(16, 17). This signal also provides a record of the enzyme’s position on its nucleic acid substrate at exquisite spatiotemporal resolution(14, 15). Put simply, SPRNT measures an enzyme’s location along a DNA strand by simply reading off the DNA sequence as it is moved through the pore, just as one would read the ticks on a ruler.

**Figure 1:**
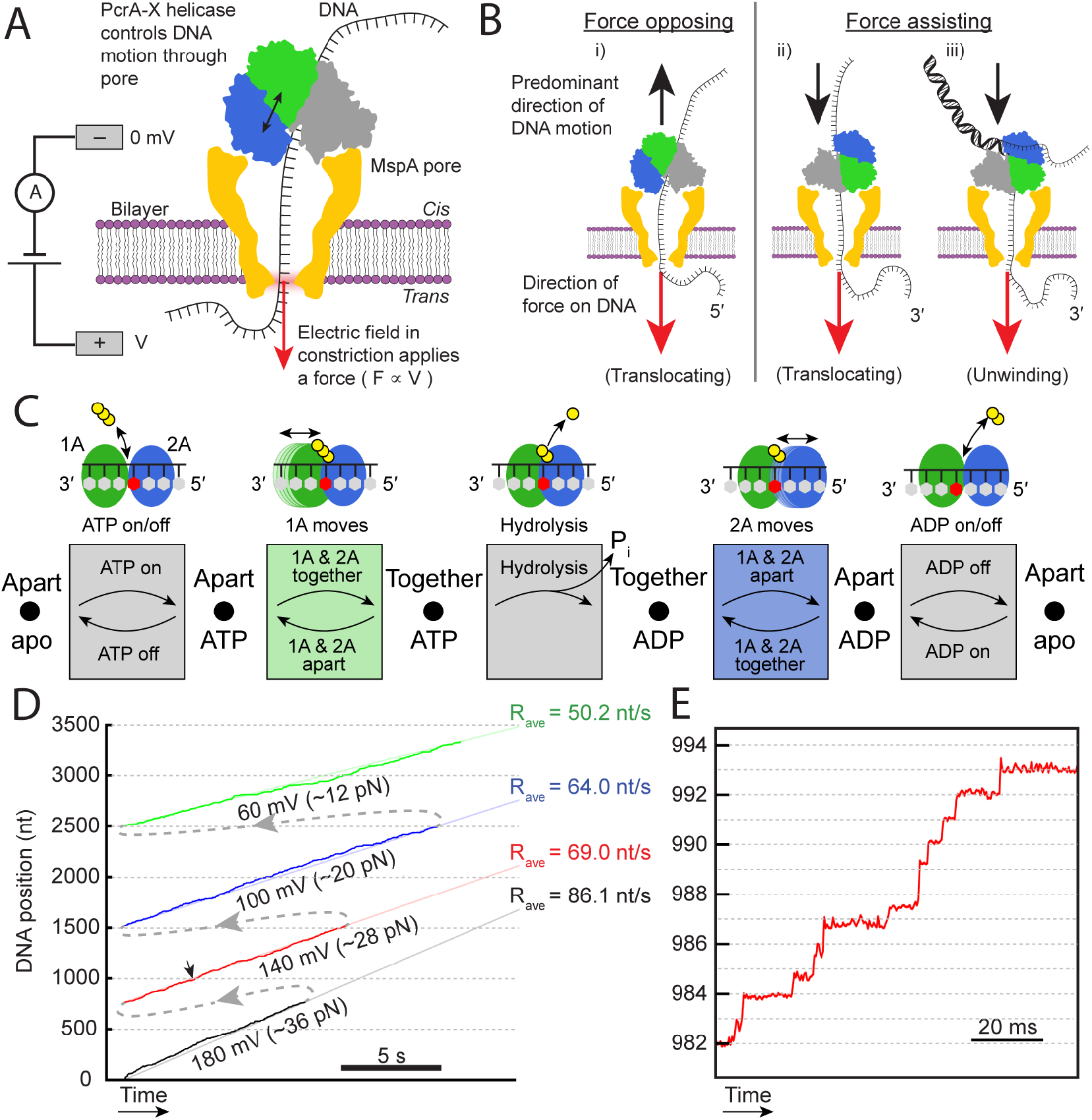
SPRNT analysis of PcrA. A) SPRNT measures the progression of DNA nucleotides through a protein nanopore. DNA bases are identified within the pore constriction by their characteristic ion current and their progression can be used to monitor the progression of a helicase along a DNA strand. RecA-like walker domains 1A and 2A are highlighted in green and blue, respectively. B) Various experimental configurations used to probe different aspects of helicase motion in force-opposing and force-assisting configurations. Force-opposing unwinding is not possible with SPRNT because the MspA constriction is only wide enough for ssDNA. C) Inchworm kinetic model of superfamily 1 and 2 helicases consists of a series of enzyme states (dots) connected by physical and chemical substeps (arrows). The substeps, in order of occurance, are 1) ATP binding/unbinding, 2) Domain 1A sliding forwards/backwards as the helicase closes/opens, respectively, 3) ATP hydrolysis, 4) Domain 2A sliding forwards/backwards as the helicase opens/closes, respectively, and 5) ADP release. In principle, each step has a forwards and backwards rate however here we treat hydrolysis as irreversible since the concentration of inorganic phosphate is effectively zero. D) Example SPRNT data, as a single PcrA helicase walks along a long repetitive strand of DNA, the applied voltage (force) is changed resulting in changes in the observed helicase stepping rate. E) Detail of D) showing easily resolved single-nucleotide steps, some as fast as 2 ms. The small black arrow in D) at ~1000 nt indicates the location of data shown in E)

SPRNT is capable of resolving sub-nucleotide nucleic acid motion on submillisecond timescales (14, 15) over distances of thousands of nucleotides while simultaneously revealing the enzyme’s sequence-specific location. SPRNT has high-throughput, enabling the collection of large datasets across a broad range of experimental parameters (18). SPRNT also applies a force to the motor enzyme which can be used to probe force-generating substeps. Previously we demonstrated SPRNT using opposing forces to enzymes translocating on ssDNA (14, 18–20) (Fig. 1Bi). Here, we expand its use to include assisting forces (Fig. 1Bii) and to, for the first time, observe duplex unwinding with SPRNT (Fig. 1Biii).

Much of what is known to date about how helicases walk along and unwind DNA comes from studies of the superfamily (SF) 1 helicase PcrA and homologs such as Rep and UvrD(6, 21–26). PcrA has two RecA-like domains, domain 1A and domain 2A, that individually bind to ssDNA(6). Between these two domains lies a highly-conserved ATP-binding pocket. PcrA translocates along the DNA lattice by progressing through a series of kinetic substates that are connected by either chemical or physical substeps(6, 22) (Fig. 1C). ATP binding induces a conformational change in the protein, bringing 1A and 2A closer together. Then, hydrolysis of ATP to ADP allows for a conformational relaxation of the protein, followed by ADP release. During the ATPase cycle, either 1A or 2A slide along the DNA backbone in alternating fashion, a so-called inchworm mechanism(6), resulting in directed motion along the DNA. Figure 1C shows a schematic of the inchworm mechanism consisting of several states, separated by chemical and physical substeps between those states. SF1 and SF2 helicases are thought to walk in similar manners because they share many core motifs for nucleic acid binding, nucleotide-triphosphate (NTP) binding, and NTP hydrolysis(2, 3, 5, 6, 27–30). While wild-type PcrA is not particularly processive in unwinding, PcrA can be activated into a highly processive helicase “PcrA-X” by locking it into its active conformation via chemical crosslinking (31) (Materials and Methods). We use this activated form of PcrA here to better understand PcrA’s mechanism of DNA unwinding and for simplicity refer to it as PcrA from here onward.

While all of this is certain, it is based on indirect lines of inference and measurement, single ATP-turnover events have yet to be directly observed at physiological conditions for any SF1 helicase in a single-molecule assay because they are too small and fast to be resolved by most single-molecule assays, other than SPRNT. Our previous SPRNT studies of the SF2 helicase Hel308 translocating along ssDNA revealed that Hel308 fed the DNA through the pore in two physical substeps per nucleotide, and revealed the chronology of the chemical substeps (ATP binding, hydrolysis, and ADP release) and conformational-change substeps that produce processive motion. In Hel308, the DNA sequence, but not the magnitude of the opposing force, strongly influenced translocation kinetics(14, 18–20). Here we have applied SPRNT to PcrA to test existing models of SF1 translocation and to determine whether the lessons learned from Hel308 can be applied broadly to SF1 and SF2 helicases. We also compare PcrA molecules translocating along ssDNA to those unwinding DNA duplexes providing insight into the mechanism of DNA unwinding.

## Results

We recorded over 600 single-molecule trajectories of PcrA helicases both unwinding and translocating along a repetitive ~6-kb-long (32) DNA construct (materials and methods, Figs. S1-S8) at different ATP and ADP concentrations and with various assisting and opposing forces. The repetitive DNA was made up of 64 repeated sections of 86 nucleotides separated by one of eight 17-base linker segments in the following configuration:

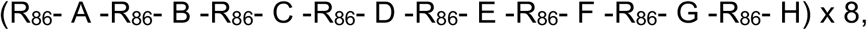

where R_86_ is the 86 base repeat section and A, B, C, etc. represent the different linker segments (full sequence in SI). This DNA strand was modified with end-adaptors that enabled use in force-opposing (translocase mode only, Fig. 1Bi) and force-assisting (translocase and unwinding mode, Fig. 1Bii, 1Biii) configurations (Figs. S1&S2, adaptor sequences in Table S1). Use of this strand regularly produced several-kb-long singlemolecule traces of individual PcrA molecules. In each full 6.6kb trace, a single PcrA molecule walks over 64 repeated instances of the 86-base repeat sequence and 8 instances of PcrA walking over each of the 8 linker sequences allowing for accumulation of significant single-molecule statistics from individual traces. In total, this dataset comprises over 50 hours of helicase-on-pore time, travelling over a combined 2.3 million bases of DNA. This, to our knowledge, represents the largest number of single-enzyme turnover steps analyzed in a single study for any motor protein (Tables S2, S3).

Because each trace encompasses thousands of individual enzyme turnover events, we were able to change the applied voltage successively during each trace to accumulate single-molecule statistics for each individual PcrA helicase at several different forces. For example, Figure 1D shows a trace in which an assisting force was changed successively from ~36 pN, to ~28 pN, to ~20 pN, and finally to ~12 pN while a single PcrA helicase translocated over a long single-stranded DNA. As force decreased, the observed mean stepping rate also decreased.

### Understanding PcrA Motion in SPRNT

In agreement with previous studies of PcrA (6, 21, 33), here we observe PcrA to take single-nucleotide steps (Figs. 1E & 1F). This is in contrast to SPRNT experiments with Hel308 helicase, which revealed two ~half-nucleotide-substeps per ATP hydrolyzed. Control experiments with other SF1 helicases Rep-X and UvrD also reveal only single-nucleotide steps (Fig S9) in our SPRNT assay. We suspect this difference arises from how PcrA, Rep, and UvrD rest on the pore rim in comparison to Hel308. Proper interpretation of our data requires that we understand which substep(s) of the inchworm mechanism results in motion of the DNA through MspA for these experiments. We previously determined that the two observed substeps per ATP hydrolyzed in Hel308 corresponded to the two conformational change substeps of the inchworm mechanism (Fig. 1C, motion of domain 1A relative to the DNA and motion of domain 2A relative to the DNA). Shifting of Hel308 on the pore rim, as opposed to a half-nucleotide motion of Hel308 along the DNA strand, is the reason for the apparent half-step motion of DNA through the pore. This suggests that one of these conformational-change substeps is responsible for the single step per ATP observed in PcrA, Rep and UvrD, while the other conformational-change substep is unobservable.

To determine whether domain 1A motion or domain 2A motion is the observable step in our experiments, we return to the schematic in Figure 1B (see also a schematic based on crystal structures of MspA and PcrA Fig. S10). In force-opposing experiments, domain 2A is in contact with the pore rim, however in force-assisting experiments, domain 1A is in contact with the pore rim. We hypothesize that different substeps are observable depending on which end of the DNA is filed through the pore, i.e. whether force is assisting or opposing PcrA motion. This is because SPRNT measures the DNA motion relative to the enzyme surface that is in contact with MspA. That is, in force-opposing experiments the observable step is motion of domain 2A backwards and forwards along the DNA (Fig. 2A); while in force-assisting experiments, the observable step is motions of domain 1A along the DNA (Figs. 2B, 2C). This is confirmed in the data that follows.

**Figure 2:**
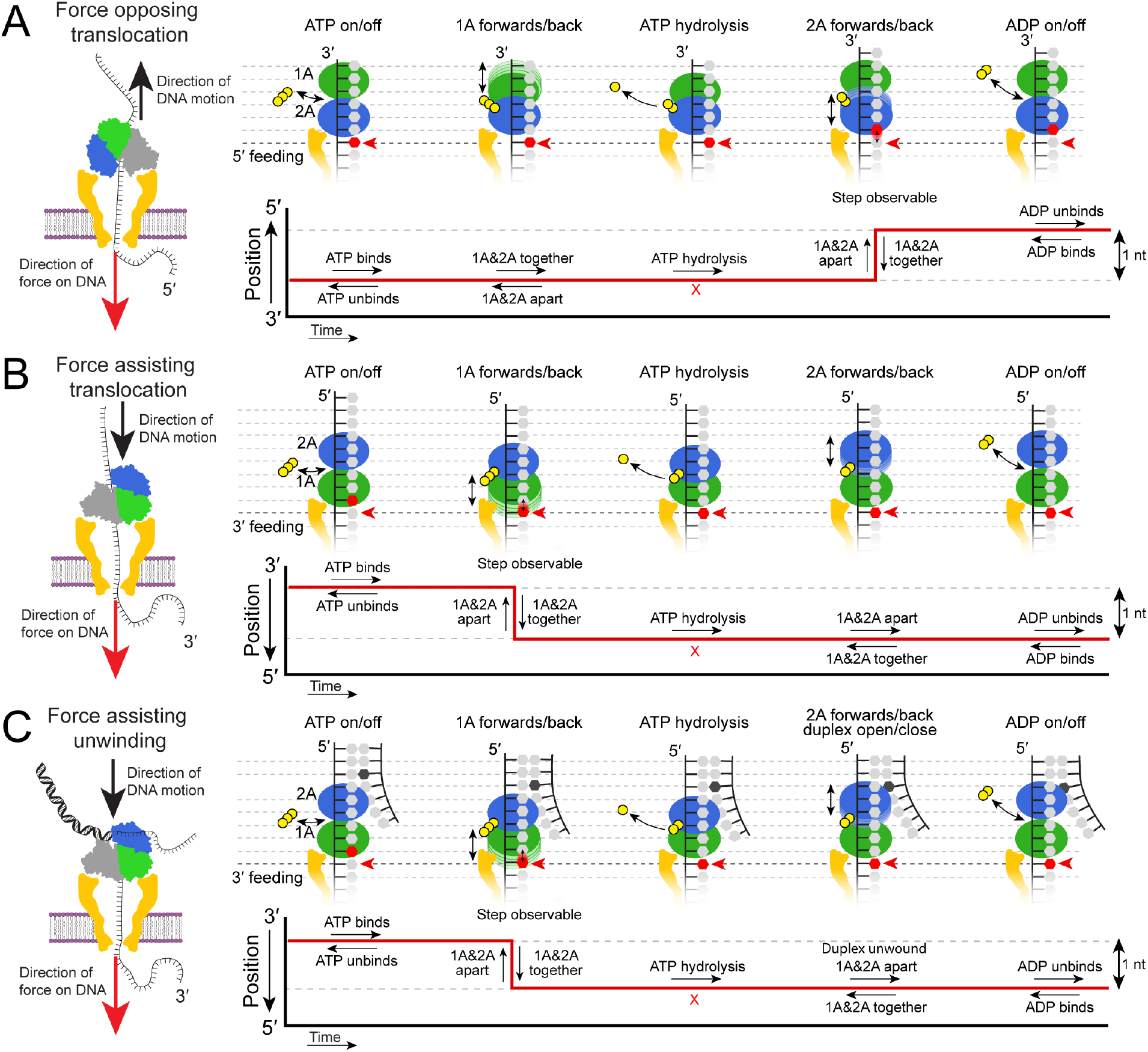
Kinetic model for PcrA translocation and unwinding in the context of SPRNT. A) *Force-opposing translocation*: Kinetic model from Figure 1A in the context of SPRNT with the 5′ end of the substrate DNA fed into the pore. The observable step occurs as the helicase transitions between “together” and “apart” and domain 2A slides along the DNA. Below in red, a schematic SPRNT data trace showing where in the ATPase cycle the observable step occurs. B) *Force-assisting translocation:* Kinetic model from Figure 1A in the context of SPRNT with the 3′ end of the substrate DNA fed into the pore. The observable step occurs as the helicase transitions between the “apart” and “together” state with domain 1A sliding along the DNA. Below in red, a schematic data trace showing where in the ATPase cycle the observable step occurs. C) *Force-assisting unwinding:* As with force-assisted translocation, the forwards/backwards motion of domain 1A is the observable step while unwinding of the complementary strand occurs either immediately prior to or coincident with domain 2A’s advance. Because of the irreversibility of the ATP hydrolysis step ([Pi] = 0), ADP is only able to directly drive backstepping in 5′ feeding experiments. High [ADP] instead increases observed step duration in force-assisting experiments. Off-pathway steps such as diffusive, [ATP]-independent movement are omitted for simplicity.

Figures 3A and 3B summarize the average speed at each force, ATP, and ADP condition. As expected, larger opposing forces slowed the motion of PcrA, while larger assisting forces sped it up; reduction in [ATP] or an increase in [ADP] lowered the average speed. Interestingly, the average speed is lower at ~12 pN *assisting* force than at ~12 pN *opposing* force, possibly due to slight distortions of the helicase that depend on how it rests on the pore rim. The range of speeds observed is quite broad; however individual enzymes have tighter speed distributions than the combined dataset (Fig. S11). In other words, there exist individual enzymes that were consistently faster or slower than average, a phenomenon known as static disorder, possibly caused by slightly different folding states or oxidative damage(34–36). For comparison to previously published measurements, Michaelis-Menten curves and inhibition curves for each applied force are shown in Figure S12. V_max_ was much lower for unwinding than for ssDNA translocation, 14 ± 3 nt/s vs. 55 ± 6 nt/s, presumably because base-pair unwinding is rate-limiting. Accordingly, saturation occurred at a lower [ATP] for unwinding experiments (K_M_ = 1.5±1.1 μM at ~28 pN assisting force) compared to ssDNA translocation (K_M_ = 6.4±1.8 μM at ~28 pN assisting force).

**Figure 3:**
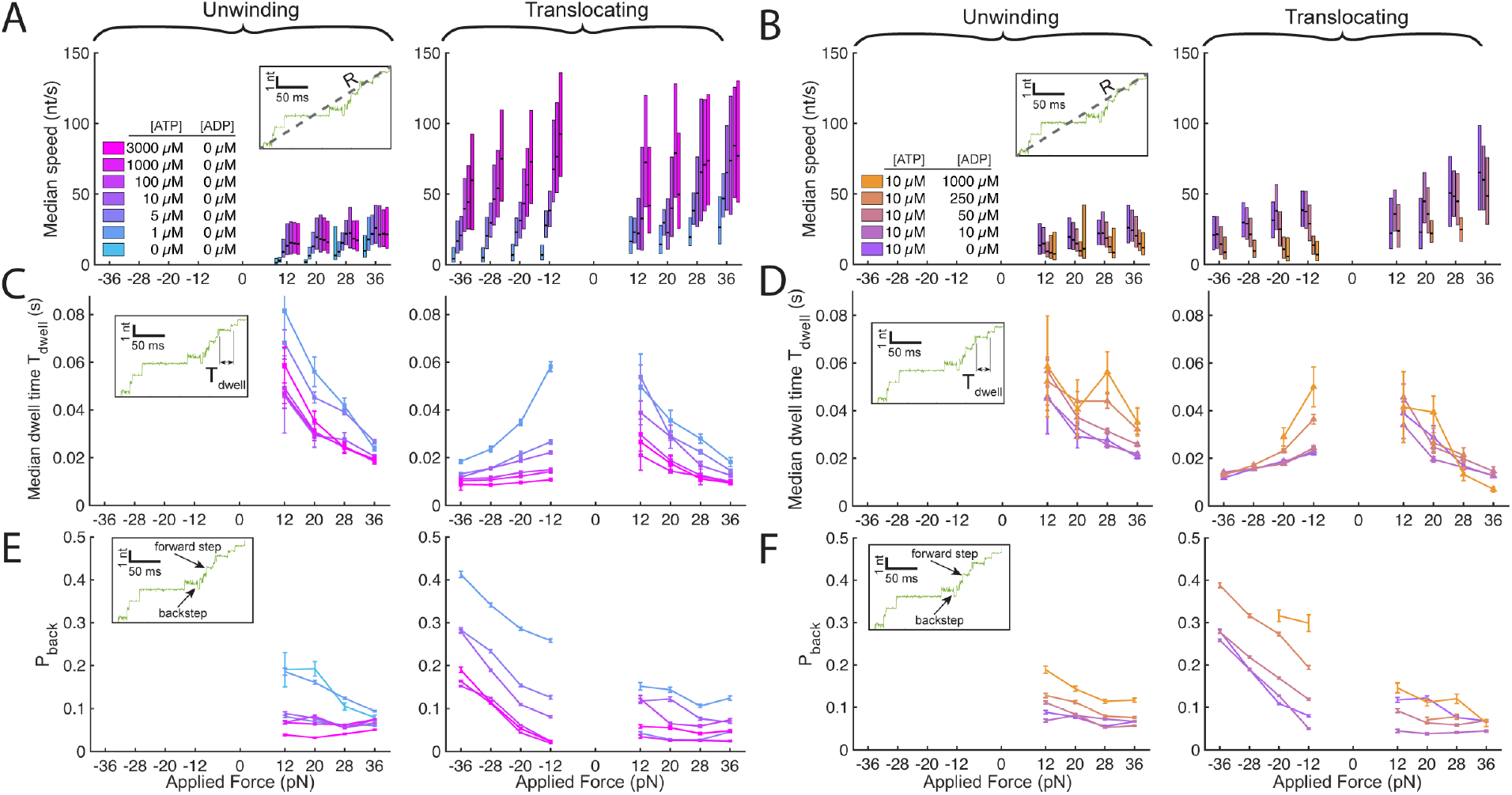
PcrA stepping kinetics at varying ATP and ADP concentrations. A) & B) Average speed for helicases in various ATP and ADP conditions at all applied forces. The speed was calculated as the average velocity taken over successive 15 nt-long stretches. The black line in the middle of each bar is the median speed while the bars in A) and B) extend to 25^th^ and 75^th^ percentiles and represent the spread in observed speed across several helicases. Positive forces are assisting, negative forces are opposing. In panel A, only the unwinding data includes an [ATP] =0 datapoint, the lowest [ATP] in the translocation data is 1μM. C) Median dwell time as a function of applied force at various ATP concentrations. D) Median dwell time as a function of force at ATP 10μM and various ADP concentrations. E) Average probability of a backward step as a function of applied force at various ATP concentrations. F) Probability of a backward step as a function of force at ATP 10 μM and various ADP concentrations. Inset graphs within each panel demonstrate features of the data being measured: speed, step dwell-time, and probability of backstepping N_back_/(N_forward_+N_back_). For the 36pN and 28pN force-opposing [ATP]=10μM, [ADP]=1000μM condition the data points were omitted since, backstepping was so prevalent (Fig. S11) that automated alignment was unreliable.

In unwinding experiments, some stepping was observed at zero [ATP] (Fig. S13). We attribute this to off-pathway, diffusive motion of the enzyme along the DNA that is rectified by the assisting force. Forwards steps with assisting force are then some combination of 1) on-pathway steps as in Figure 2 and 2) force-driven, off-pathway steps. We account for this effect with a modified Michalis-Menten equation of the form:

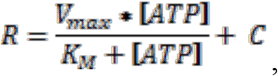

which includes the normal Michaelis-Menten term with the addition of an [ATP]-independent constant C, where C is determined by the stepping rate at [ATP]=0 (Fig. S12). Put simply, the total rate R, is the sum of the on-pathway rate and off-pathway rate. The ratio C/R yields what fraction of the observed steps are off-pathway at each applied force. Intriguingly, during unwinding, only C increases with force, while the V_max_, which is associated with ATP-hydrolysis-driven motion, remains constant (Fig. S12). This suggests that only off-pathway steps are affected by the assisting force while on-pathway ATP-hydrolysis-driven unwinding is not affected by force. This is because force applied during SPRNT couples only to steps that result in motion of DNA through the pore, since the applied force on the DNA can only do work on the system when the force results in a change in DNA position(37). This is consistent with the model presented above. In force-opposing experiments, force modifies the forwards and backwards rates of domain 2A along DNA (Fig. 2A), while in force-assisting experiments, force affects the forwards and backwards rates of domain 1A (Figs. 2B, 2C). Because unwinding is significantly slower than translocation, it is reasonable to suppose that the rate-limiting substep is the one in which duplex unwinding occurs. Because domain 2A advance is rate-limiting, force-driven changes in domain 1A motion should have little effect on the rate of on-pathway unwinding.

SPRNT’s ability to resolve single-nucleotide steps enables us to see the underlying kinetic details such as dwell-time (Figs. 3C, D) and backstepping probability (Figs. 3E, F) and to observe how these observables give rise to average behavior. For example, as the opposing force was increased, the median dwell-time at each nucleotide step (Tdwell) during ssDNA translocation decreased. On its own, this is surprising as one might expect the opposite since the average speed decreased with opposing force. This apparent paradox is resolved by noting that the probability of a backstep (P_back_) increased significantly with opposing force. Thus, opposing force reduces PcrA’s average speed by increasing its propensity for backstepping rather than simply reducing the forwards stepping rate.

As expected, reduced [ATP] led to increased dwell-times at all applied forces. Surprisingly, reduced [ATP] also led to increased P_back_ at all applied forces. The inchworm mechanism does not predict that insufficient ATP will cause backwards steps, so we attribute this motion to off-pathway behavior, consistent with diffusive motion along the DNA as is seen with higher assisting forces. Such diffusion may generally occur at a low rate and only become visible when low ATP or high opposing force sufficiently inhibit on-pathway behavior.

To understand the effect of [ADP] on PcrA kinetics we held [ATP] at 10μM while titrating [ADP] up to 1000 μM. As [ADP] was increased, the average speed reduced linearly with [ADP], consistent with competitive inhibition of ATP-binding. At the level of individual enzyme steps, force-assisting and force-opposing data are remarkably different. In force-opposing experiments, increased [ADP] causes a significant increase in backstepping. At higher opposing forces (≥28 pN) and [ADP] = 1000μM, backstepping was so prevalent as to render alignment to the DNA sequence impossible (Fig. S14). In contrast, in force-assisting experiments little change in P_back_ was observed with changing ADP concentration. This discrepancy can be understood by looking at the model in Figure 2. The presence of ADP should allow for backstepping of domain 2A which is observable during force-opposing experiments. However, because the concentration of inorganic phosphate is essentially zero, hydrolysis is effectively irreversible. Therefore, backstepping of domain 1A, which is the observable step in force-assisting experiments, is unaffected by the presence of ADP. While assisting force measurements are essentially blind to forwards and backwards motions of domain 2A, these backsteps should instead manifest in the data as longer dwell-times, as is apparent in the unwinding data of Figure 3D. Step dwell-times for force-assisting translocation do not appear to be significantly longer at high [ADP] suggesting ADP-driven backstepping of domain 2A requires a force opposing PcrA’s motion such as the nanopore rim in force-opposing experiments or the DNA duplex in force-assisting unwinding experiments. Extrapolating P_back_ to zero opposing force suggests domain 2A backsteps little at low opposing forces.

This insight also sheds light on the kinetics of PcrA during unwinding and suggests that opposing force applied during SPRNT is analogous to the opposing force applied by a DNA duplex during unwinding. During force-opposing translocation experiments, numerous backsteps are observed especially with larger opposing forces. By analogy, we hypothesize that during unwinding, domain 2A makes several attempts to invade the DNA duplex before successfully stepping forward. More attempts are made if the energy barrier is high, either via higher opposing force or a stronger DNA duplex. The unbinding of ADP ultimately rectifies the forwards step of domain 2A. Comparison of conditional dwell-time distributions further support this interpretation (Fig. S15).

### DNA Sequence Affects both Translocation and Unwinding PcrA Kinetics

Recent studies of SF1 helicase UvrD(38) and SF2 helicase Hel308(14, 18, 19) have shown that ssDNA translocation speed can be strongly affected by the underlying DNA base composition. SPRNT is well-suited to detection of sequence-dependent kinetics because SPRNT measures enzyme position by detecting the DNA sequence directly (Methods), thereby revealing an enzyme’s sequence-specific position along the DNA strand. The long, repetitive DNA strand used in this study allows for collection of data in which a single PcrA enzyme walks over the same 86-base long DNA sequence multiple times at several different applied forces allowing for accumulation of significant sequence-specific kinetic data. Figure 4A shows the median dwell-time for a forward step as a function of location within the 86-base-long repeat section of the antisense strand for both dsDNA unwinding and ssDNA translocation, T_Unwind_ and T_Trans_, respectively (sense strand in Fig. S16). Average dwell-times vary by an order of magnitude depending upon where PcrA is along the sequence for both unwinding and translocation.

**Figure 4:**
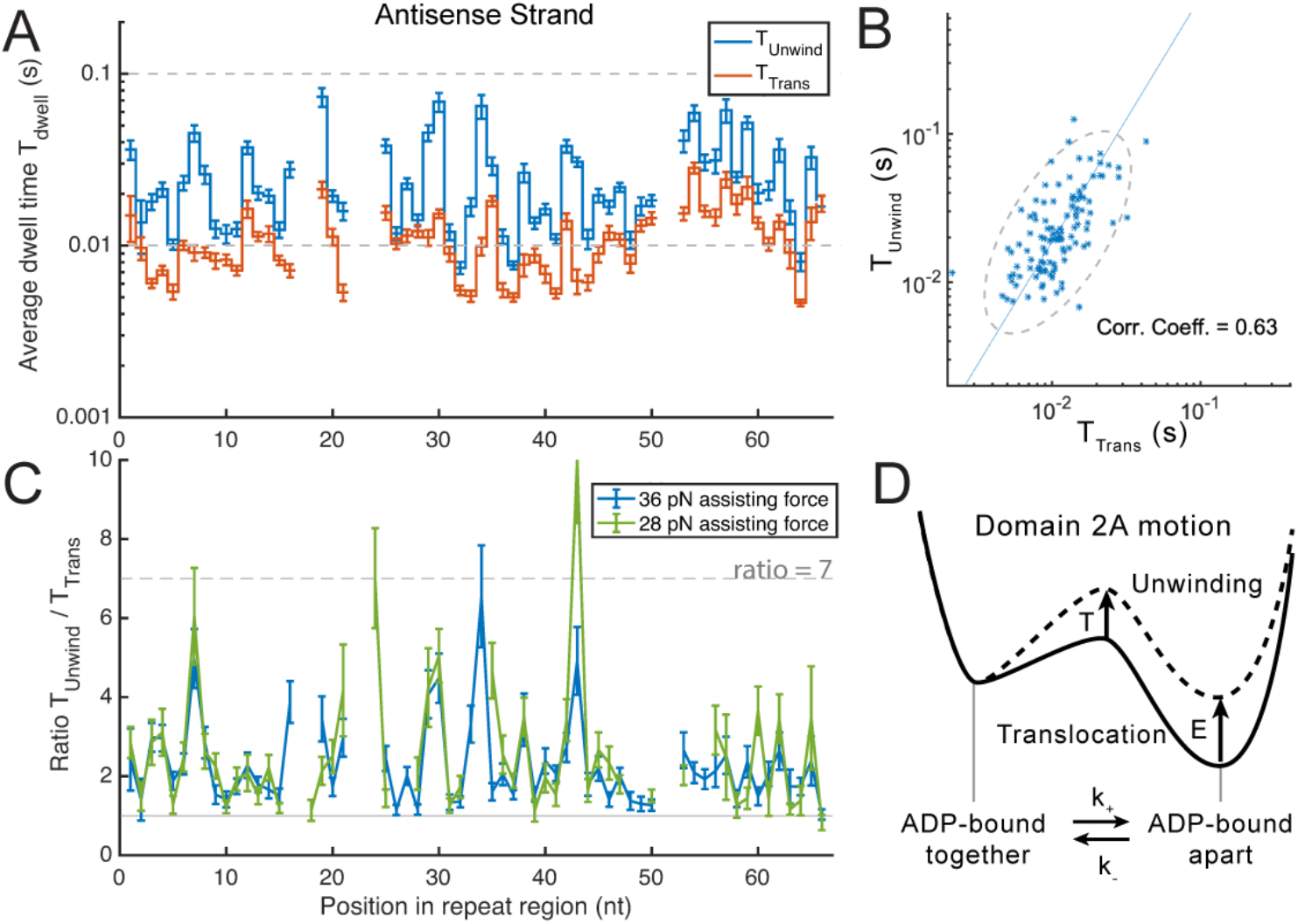
A) Average dwell time for a forwards step for PcrA walking along the antisense strand during dsDNA unwinding (blue) and ssDNA translocation (orange) with 36 pN assisting force. The variation in dwell times as a function of position along the DNA indicates strong sequence dependence during translocation and unwinding. B) Scatter plot of ssDNA translocation dwell time T_Trans_ vs. unwinding dwell time T_Unwind_ for both sense and antisense strand data on a log/log scale. C) The ratio of T_Unwind_ to T_Trans_ as a function of location within the repeat section of the DNA antisense strand. The ratio in step dwell times is sequence dependent and consistent across different applied forces. Gaps within the data are in locations in which the ion-current trace has low signal-to-noise and single nucleotide enzyme steps cannot be easily resolved. D) Transition state theory energy diagram showing how duplex unwinding modifies the forwards and backwards rates of domain 2A motion in the model in Figure 2.

By leveraging the underlying structure of our DNA substrate, we were able to determine that the variation in dwell-time was due to different bases within the PcrA. In SPRNT, DNA bases are read at the pore constriction which is separated by ~12 nucleotides from motor enzyme. During force-assisting experiments PcrA enters the linker region several nucleotides before the linker region enters the pore constriction. This separation serves as a natural experiment in which we can compare step dwell-times while MspA passes over the tail end of the repeat section while PcrA walks over different linker sequences (Fig. S17). Significant sequence-dependent effects begin ~15-20 nucleotides prior to the linker’s entry into the pore constriction. This is similar to previous SPRNT results with Hel308 helicase in which sequence-dependent effects were found to be due to two sites within Hel308 separated 17 and 20 nucleotides from the pore constriction (19).

While one might assume the work required to unwind DNA would dictate the rate of unwinding, our data indicate that the ssDNA translocation rate plays an important role in unwinding. Even though unwinding is considerably slower than translocation, at each location along the DNA strand T_Unwind_ and T_Trans_ are correlated (Fig. 4B, Fig. S16) suggesting that the rate-limiting reaction substep of DNA unwinding is coincident with a rate-limiting substep for translocation on ssDNA. In other words, unwinding modifies the rate constants of forwards and backwards motions of domain 2A in Figure 2, rather than adding a new rate-limiting substep to the cycle.

The ratio of T_Unwind_ to T_Trans_ is often used to compare the difference between translocation and unwinding(5, 39, 40). If the ssDNA sequence alone fully determined the rate of DNA unwinding, we would expect this ratio to be flat over the 86-base-long region. Instead the ratio of T_Unwind_ to T_Trans_ also depends on the underlying sequence (Fig. 4C) implying that duplex identity also plays a role in determining sequence kinetics. Interestingly, the ratio of T_Unwind_ to T_Trans_ at 36 pN and at 28 pN assisting force shows similar sequence dependent patterns (Fig. 4C); and the same pattern of sequence dependence is also apparent at 20 pN and 12 pN (Fig. S18). This supports our conclusion that the assisting force has no effect on domain 2A motion in which unwinding occurs.

Figure 4D shows a transition state theory interpretation of this result for the substep in which domain 2A moves. In this model, the free energies of the bound states and transition state of the energy-diagram are determined by the ssDNA sequence (Fig. 4D, solid line). Opposing force, be it DNA duplex or applied by SPRNT, modifies the energy landscape by increasing the transition state energy by some energy T and the bound-state energy by some energy E (Fig. 4D, dashed line). This approach suggests a more complicated relationship exists between T_Unwind_ and T_Trans_ than simply T_Unwind_ ∝ T_Trans_ (Fig. S16) because both the forwards and backwards rates of domain 2A motion are modified by force.

## Discussion

SPRNT allows for direct observation of individual ATPase cycles of helicases, revealing fundamental mechanochemical substeps. By changing experimental conditions such as [ATP] and [ADP], and by applying assisting or opposing forces we probed substeps of the inchworm-mechanism to build a detailed picture of how PcrA unwinds DNA. As PcrA walks along ssDNA, the presence of a complementary DNA duplex exerts an opposing force which counters the advance of domain 2A. Larger opposing forces increase the backstepping rate of domain 2A, impeding forwards progress. This backstepping is also ADP-dependent, thus, unbinding of ADP helps to rectify PcrA’s progress against an opposing force. SPRNT’s ability to associate PcrA’s steps with the underlying DNA sequence reveals a complex mechanochemical landscape for duplex unwinding that is dependent upon both the ssDNA sequence and the duplex stability. In this sense, PcrA is like a snowplow: the amount of snow in front of the plow is important but so too is the amount of traction available to the wheels. Differences in base-pair stability vary how much work is required to unwind the DNA duplex while the underlying ssDNA sequence significantly affects how quickly PcrA can perform that work. Sequence-dependent translocation kinetics should be investigated as a source of sequence-dependent unwinding behavior in other enzymes.

Our data show that the individual motion of both RecA domains, off-pathway stepping, and effects of DNA sequence on translocation and unwinding are all important details of PcrA kinetics. These new details suggest that a full description of helicase behavior is far richer than models commonly used to describe helicase motion. Rather than existing on a spectrum from passive to active(5, 39, 40) PcrA exhibits a range of “activity” depending on both the ssDNA sequence and nucleotides being unwound.

The large variation in sequence-dependent stepping is significant when considering bulk average kinetic data such as average speed. The order-of-magnitude variation in step dwell-times means that just a few steps that are slow because of their sequence contexts end up dominating PcrA’s average behavior. This, in addition to static disorder, helps to explain the broad range of speeds measured. Characterizing PcrA by its average behavior misses important features that our SPRNT data are able to reveal. Our data cover a broad range of sequence contexts (Figs. S3-S5) suggesting that PcrA kinetics depend on the underlying sequence in general and not just for one particular DNA sequence motif. Comparison of the T_Unwind_/T_Trans_ ratio with the base pairing binding energies of the underlying sequence does not reveal a simple relationship between a single base-pair and the effect of the DNA duplex on unwinding speed (Fig. S19). The crystal structure of PcrA reveals numerous contacts between the DNA duplex and the exterior of PcrA, suggesting that several DNA bases within the duplex may be important for determining translocation and unwinding speed. Systematic studies on a far broader range of sequence contexts will be needed to fully understand the cause of such sequence-dependence.

The fact that sequence-dependent translocation has now been observed in three different superfamily 1 and superfamily 2 helicases (PcrA, Hel308(19), and UvrD(38)) suggests that sequence-dependence is a common behavior of helicases. Unlike protein filaments (e.g. actin), DNA is not a homogeneous track; sequence-dependent behavior may be the norm rather than the exception. Strong sequence-dependent enzyme kinetics such as those observed in our data likely affect PcrA’s role *in vivo* and could thereby exert selective pressure on both DNA and protein evolution. Therefore, sequence-dependent behavior should be carefully considered in future studies of any enzyme that walks along DNA or RNA since the sequence-dependent kinetics may reveal essential features of an enzyme’s function. Such effects are almost certainly used by life to achieve various ends; and SPRNT is well suited to discovering how and why such sequence dependence occurs and opens the possibility of uncovering enzyme functions that were hereto unknown.

## Materials and Methods

A single MspA pore is prepared in an unsupported phospholipid bilayer as previously reported (15). The pore is prepared with asymmetric salt conditions to provide a low-salt environment for PcrA while still ensuring a strong ion-current signal. The *trans* well contains 500mM KCl, 10mM HEPES, buffered to pH 8.00±0.05 and the *cis* well contains 100mM KCl, 10mM MgCl_2_, 10mM HEPES, various [ATP] and [ADP], buffered to pH 8.00±0.05. Unless otherwise noted, all experiments occur at room temperature. A voltage is applied across the membrane by an Axopatch 200B patch-clamp amplifier (Molecular Devices). Once a ss nucleic acid end is captured by the electric field in the pore, it is rapidly pulled through the pore until the bound motor enzyme contacts the rim of the nanopore. From this moment onward, the enzyme controls the motion of the nucleic acid through the pore, until the enzyme falls off the nucleic acid.

Data is acquired at a sampling rate of 50 kHz filtered with an 8-pole 10 kHz Bessel-filter. Data is then digitally down-sampled to 5 kHz for analysis using a boxcar-filter. Single-enzyme traces are detected automatically via thresholding. Reads begin when current drops below 80% of the open pore current and end when ion-current returns to above 94% of open pore current for more than 10 datapoints. Putative events are then evaluated by-hand by recognition of ion-current patterns corresponding to sense or antisense strand and 3′ and 5′ feeding (Figs. S3, S4, S5). 5′ feeding antisense data is not used in this study because this DNA sequence produces a long region of low-current contrast which interferes with detection of individual enzyme steps. Steps in ion current data are detected using a change-point algorithm previously described (14, 18). Ion current values are then automatically aligned to the corresponding ion-current consensus (Figs. S3, S4, S5) using a previously described dynamic-programming alignment algorithm (17).

Varying forces are applied by changing the voltage applied across the membrane. DNA-enzyme complexes were captured at 180mV and then switched at intervals to 60 mV, 100 mV, 140 mV. Current estimates of applied force are 0.20 ± 0.04 pN/mV based on how DNA stretches within the pore at different applied forces(41). Applied voltages of 60 mV, 100 mV, 140 mV, and 180 mV correspond to 12 ± 2 pN, 20 ± 4 pN, 28 ± 6 pN, and 36 ± 7 pN. In comparison to the literature, these forces appear quite large, but the way in which the force is applied is key to how the applied force affects enzyme stepping kinetics(12). All experiments were performed at room temperature.

Wild-type PcrA is a poor helicase, only unwinding a few bases at a time, however recent studies revealed that by constraining the conformation of PcrA’s accessory domains into a “closed” configuration, PcrA could be transformed into a so-called “superhelicase” capable of unwinding thousands of bases at a time against large opposing forces(31). In this study, we use this activated PcrA helicase from *Bacillus stearothermophilus* (accession no. P56255) with the mutations (C96A, C247A, N187C and L409C) “PcrA-X.” The use of chemically activated PcrA in this study enabled us to observe processive unwinding by a helicase using SPRNT for the first time. Without crosslinking into the closed form, these monomeric enzymes switch strands after unwinding just a few base pairs, and translocate on the opposite strand while in the open form, letting the DNA rezip(42). This behavior plays a regulatory role by preventing unwinding of a long stretch of DNA until a partner protein comes and stabilizes the closed form(31). Ideally, crosslinking simply disables strand switching, thereby providing insight into the mechanism of processive unwinding. As noted in Arslan *et al*., the chemical crosslinking that locks PcrA into the ‘closed’ state required the removal of two natural cysteines (C96 and C247) which decreased ATPase activity when compared to WT PcrA. The crosslinking reaction itself does not seem to change ssDNA translocation because control experiments of this PcrA mutant before chemical crosslinking showed similar translocation behavior (Fig. S20).

PcrA helicase was purified and crosslinked as previously described (Arslan, et al., 2015). Briefly, we inserted PcrA sequence between NdeI and BamHI sites of pET-11b vector and transformed it into E. coli BL21(DE3) cells grown in TB medium. When optical density reached OD_600_ = 0.6, the protein overexpression was induced with 0.5 mM IPTG. The cells were incubated overnight at 18°C and harvested by centrifugation at 10,000 x g. Our PcrA contains 6x-His tag on its N-terminus for Ni-NTA affinity-column-based purification. The cell pellet was resuspended in a lysis buffer and cells were lysed with a sonicator. After binding PcrA to Ni-NTA column and several washes, PcrA was eluted with 150 mM imidazole-containing buffer. PcrA concentration was kept below 4 mg/mL (≈50 μM) to avoid aggregation.

Our PcrA mutant has two native cysteines C96A and C247A removed, while N187C and L409C are introduced for crosslinking. New C187 and C409 are crosslinked with BM(PEG)2 (1,8-bismaleimido-diethyleneglycol) to lock PcrA in the closed conformation and form PcrA-X. Optimal crosslinking condition is achieved at PcrA concentration between 20 and 25 μM, with PcrA to BM(PEG) ratio of 1:5 (Fig. S21).

The long, repetitive DNA strand was replicated within *E. coli* (DH5-alpha) in a pET28a plasmid vector. Original assembly of the repetitive DNA was as described previously(32). After purification using a Qiagen Maxiprep plasmid purification kit (Qiagen). The 64x repeat was then cleaved out of the plasmid using XbaI, and HindIII. Simultaneously the remainder pET28a plasmid was digested with DdeI and Fnu4HI producing numerous small DNA segments and one 6 kb-long repetitive target DNA. The 6 kb long 64x repeat was then purified using SPRIselect beads (Beckman Coulter) at a 1 to 0.4 v/v ratio. Adapters (Fig. S2) were pre-annealed and then ligated to the purified repeat DNA using T4 ligase.

Instantaneous enzyme speed in nt/s is measured using the time to travel 15 nucleotides in consecutive increments yielding several speed measurements for each single-molecule trace. Unless otherwise noted, step dwell-times are for forwards steps that happened after another forwards step. This conditional step characterization was taken into account to ensure full ATP turnover for each step. In the model of Figure 2, a forwards step that follows a backstep would not have the same dwell-time distribution as forwards steps that follow a backwards step since a step’s initial conditions modify which states the enzyme passes through before finalizing the step (18). A “backstep” refers to backwards steps occurred immediately after forwards steps; consecutive backsteps were rare. See the Supplement for further discussion.

Rep-X helicase was prepared as described previously(31). *E. coli* UvrD helicase was acquired from Quidel, (San Diego, USA).

## Data Availability

The Matlab workspace including all data for this manuscript is uploaded to figshare and is available for reviewers at the following link: https://figshare.com/s/6f53a9f1cff8d5814fa1.

## Funding

This work was supported by the National Institutes of Health National Human Genome Research Institute [R01HG005115] and the National Institute of General Medical Sciences [R35 GM 122569]. M.G. was supported by the NSERC Canada PDF.

## Author contributions

A.H.L., J.M.C., T.H., and J.H.G. designed the research. J.R.H., H.C.K., S.J.A. M.d.S., J.W.M., J.L.B., K.S.B., and H.H. acquired the data. A.H.L. analyzed the data. R.T., M.G. and D.B. produced PcrA-X helicase. A.H.L., J.M.C., M.G., T.H., and J.H.G. wrote the paper.

## Notes

https://figshare.com/s/6f53a9f1cff8d5814fa1

### Competing Interest Statement

The University of Washington has filed a provisional patent on the SPRNT technology.

