## Supplementary material for "Sequence-dependent mechanochemical coupling of helicase translocation and unwinding at single-nucleotide resolution": Fig. S

Supplement for **Sequence-dependent coupling of helicase translocation and unwinding**

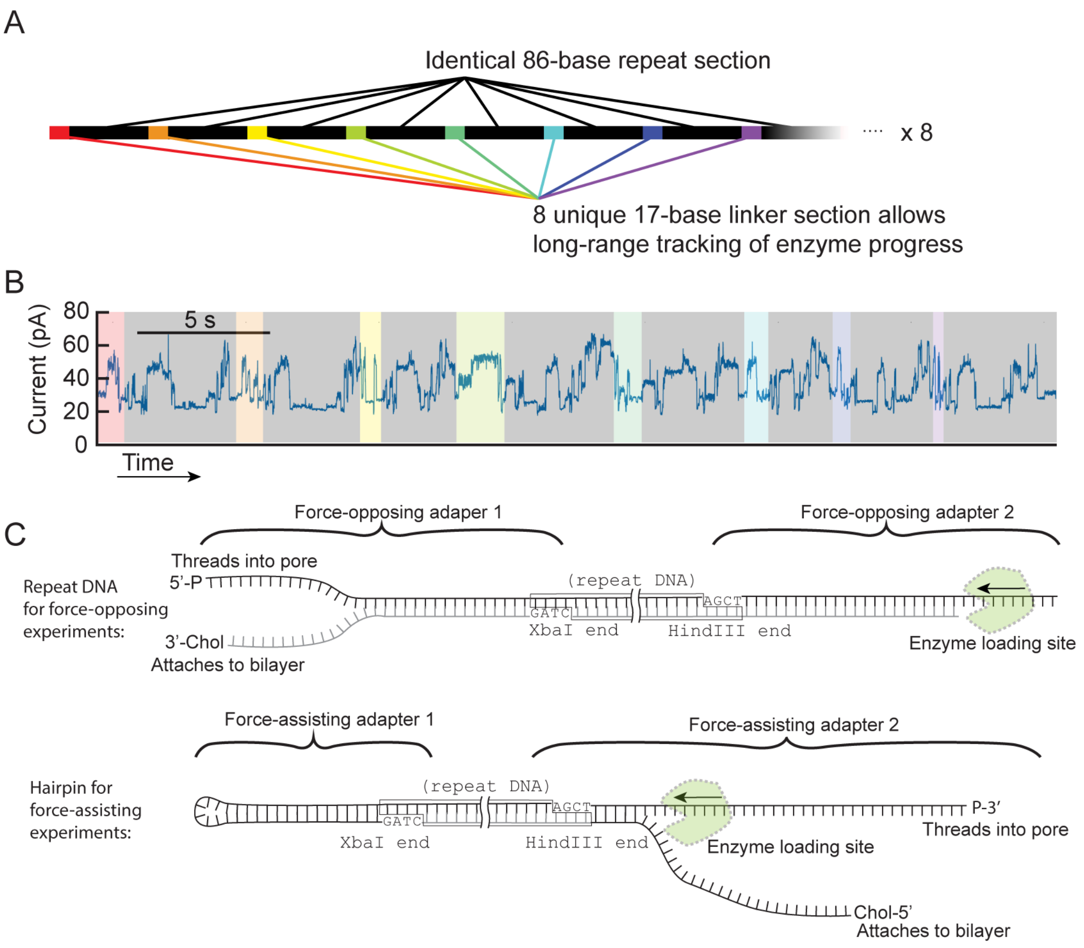

**Figure S1 | Construction repetitive of DNA test-track.** A) The DNA strand used in this experiment consisted of a repeating 86 nt section ($R_{86}$) of bases linked together by 8 unique 17-base long linker sequences in the following configuration[1]:

$$(R_{86} – A – R_{86} –B – R_{86} – C – R_{86} – D – R_{86} – E – R_{86} – F – R_{86} – G – R_{86} – H –) \times 8$$

for a total of 64 repeats. The sequence of this 8x repeat is shown below. B) Nanopore reads of the sense and antisense strands of this construct reveal repetitive sequence interspersed with unique sequences for each of the eight linkers. This sequence was inserted into a Pet28a plasmid is described previously[1] between the XbaI and HindIII cut sites. This allowed for restriction digestion and ligation of adapters to the left and right ends of the repeat sequence. Briefly, the repeat-containing-plasmid was digested with XbaI, HindIII, DdeI, and Fnu4HI (New England Biolabs) leaving a single, 6kb-long repetitive strand and many smaller plasmid fragments. The 6kb-long repeat DNA was then purified using SpriSelect beads (Beckman Coulter) at a ratio of 0.4:1 bead volume to sample volume. Adapters such as those shown in C) were then ligated onto the XbaI or HindIII sides of the repeat DNA to enable force-assisting or force-opposing reads. XbaI and HindIII versions of all four adapters shown (8 adapters total) were used to enable all possible configurations for both sense and antisense reads. Sequences for each adapter and the full 64x repeat are provided below.

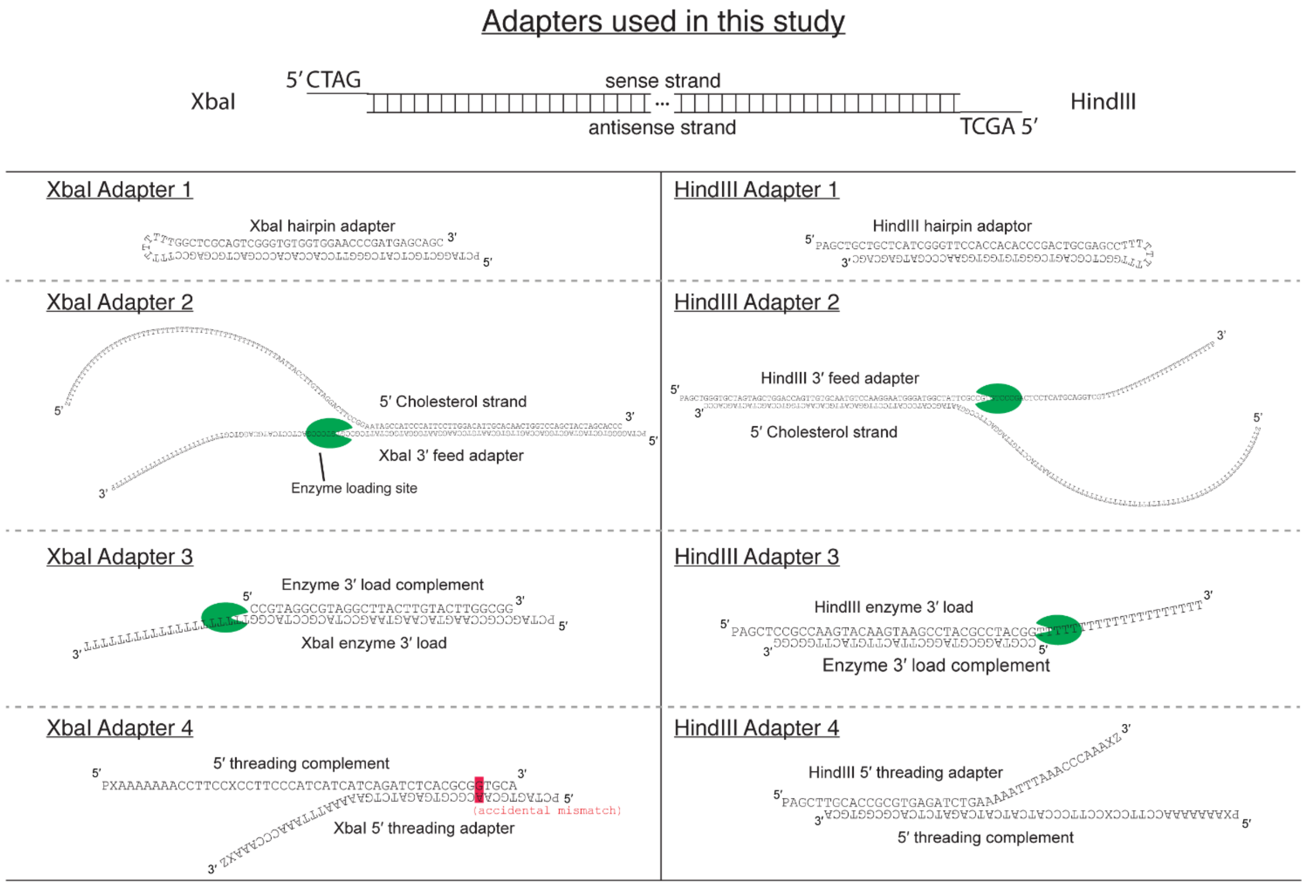

P = phosphate, Z = Cholesterol, X = abasic

Recipes:

1) to get 5’ threading sense reads:.....................................................................attach XbaI adapter 4 + HindIII adapter 3
2) to get 5’ threading antisense reads:...............................................................attach XbaI adapter 3 + HindIII adapter 4
3) to get 3’ threading hel mode asense reads and trans mode sense.............attach XbaI adapter 2 + HindIII adapter 1
4) to get 3’ threading hel mode sense reads and trans mode asense..............attach XbaI adapter 1 + HindIII adapter 2
5) to get 3’ threading hel mode sense and asense reads:................................attach XbaI adapter 2 + HindIII adapter 2
6) to get 5’ threading sense and antisense reads:............................................attach XbaI adapter 3 + HindIII adapter 3

**Figure S2 | DNA adapters used in this study** (PAN oligo synthesis, Stanford).

### Table S1 | DNA adapter sequences

| | |
|---|---|
| Type 1 adapters | Prevents pore feeding and enables transition from helicase to translocase mode |
| XbaI hairpin adapter | PCTAGGCTGCTCATCGGGTTCCACCACACCCGACTGCGAGCCTTTTTTTTTGGCTCGCAGTCGGGTGTGGTGGAACCCGATGAGCAGC |
| HindIII hairpin adapter | PAGCTGCTGCTCATCGGGTTCCACCACACCCGACTGCGAGCCTTTTTTTTTGGCTCGCAGTCGGGTGTGGTGGAACCCGATGAGCAGC |
| Type 2 adapters | Encourages force-assisted helicase mode events |
| 5’ cholesterol strand (z = chol) | ZTTTTTTTTTTTTTTTTTTTTTTTTTTTTTTTTTTTTTTTTTTTTTTTTTAATTACCTTGTTAGGACTTCCGGAATAGCCATCCCATTCCTTGGACATTGCACAACTGGTCCAGCTACTAGCACCC |
| XbaI 3’ feed adapter | PCTAGGGGTGCTAGTAGCTGGACCAGTTGTGCAATGTCCAAGGAATGGGATGGCTATTCGCCGTGTCCCGACTCCTCATGCAGGTCGTTTTTTTTTTTTTTTTTTTTTTTTP |
| HindIII 3’ feed adapter | PAGCTGGGTGCTAGTAGCTGGACCAGTTGTGCAATGTCCAAGGAATGGGATGGCTATTCGCCGTGTCCCGACTCCTCATGCAGGTCGTTTTTTTTTTTTTTTTTTTTTTTTP |
| Type 3 adapters | For force-opposing translocase mode events |
| Enzyme 3’ load complement | CCGTAGGCGTAGGCTTACTTGTACTTGGCGG |
| XbaI enzyme 3’ load adapter | PCTAGCCGCCAAGTACAAGTAAGCCTACGCCTACGGTTTTTTTTTTTTTTTTTTTT |
| HindIII enzyme 3’ loading adapter | PAGCTCCGCCAAGTACAAGTAAGCCTACGCCTACGGTTTTTTTTTTTTTTTTTTTT |
| Type 4 adapters | Promotes 5’ feeding for force-opposing translocase experiments |
| 5’ threading complement | PXAAAAAAAACCTTCCXCCTTCCCATCATCATCAGATCTCACGCGGTGCA |
| XbaI 5’ threading adapter | PCTAGTGCAACGCGTGAGATCTGAAAAAATTTAAACCCAAAXZ |
| HindIII 5’ threading adapter | PAGCTTGCACCGCGTGAGATCTGAAAAAATTTAAACCCAAAXZ |

**Sequence of 64x repeat sequence** including XbaI and HindIII cutsites at the 5′ and 3′ ends respectively (assembly described previously[1]). Lower case letters indicate the location of linker regions while upper case letters indicate the repeat sequence. Every eighth repeat is delineated with a line indicating the location at which the 8-linker sequence repeats. This sequence is essentially a nano-scale ruler for DNA position, in which the individual bases are the minor ticks and the 8 unique linkers are the major ticks.

```
TCTAGAtagacggcatgGGGAGACGCGACCGAAATGGTGAAGGACGGGTCCAGTGCTTCGGCACTGTTGAGTAGAGTGTGAGCTCCGTAACTGGTCGCGTCAC
gtaagatgctccggttaGGGAGACGCGACCGAAATGGTGAAGGACGGGTCCAGTGCTTCGGCACTGTTGAGTAGAGTGTGAGCTCCGTAACTGGTCGCGTCAC
tgatgtaccgttgagcaGGGAGACGCGACCGAAATGGTGAAGGACGGGTCCAGTGCTTCGGCACTGTTGAGTAGAGTGTGAGCTCCGTAACTGGTCGCGTCAC
tcgctagagcatggtttGGGAGACGCGACCGAAATGGTGAAGGACGGGTCCAGTGCTTCGGCACTGTTGAGTAGAGTGTGAGCTCCGTAACTGGTCGCGTCAC
tggggcacgccgtctggGGGAGACGCGACCGAAATGGTGAAGGACGGGTCCAGTGCTTCGGCACTGTTGAGTAGAGTGTGAGCTCCGTAACTGGTCGCGTCAC
ttactgcgaccgcaataGGGAGACGCGACCGAAATGGTGAAGGACGGGTCCAGTGCTTCGGCACTGTTGAGTAGAGTGTGAGCTCCGTAACTGGTCGCGTCAC
gcgcgcaaccgggtagaGGGAGACGCGACCGAAATGGTGAAGGACGGGTCCAGTGCTTCGGCACTGTTGAGTAGAGTGTGAGCTCCGTAACTGGTCGCGTCAC
gtaactcacggcgctatGGGAGACGCGACCGAAATGGTGAAGGACGGGTCCAGTGCTTCGGCACTGTTGAGTAGAGTGTGAGCTCCGTAACTGGTCGCGTCAC
gctagatagacggcatgGGGAGACGCGACCGAAATGGTGAAGGACGGGTCCAGTGCTTCGGCACTGTTGAGTAGAGTGTGAGCTCCGTAACTGGTCGCGTCAC
gtaagatgctccggttaGGGAGACGCGACCGAAATGGTGAAGGACGGGTCCAGTGCTTCGGCACTGTTGAGTAGAGTGTGAGCTCCGTAACTGGTCGCGTCAC
tgatgtaccgttgagcaGGGAGACGCGACCGAAATGGTGAAGGACGGGTCCAGTGCTTCGGCACTGTTGAGTAGAGTGTGAGCTCCGTAACTGGTCGCGTCAC
tcgctagagcatggtttGGGAGACGCGACCGAAATGGTGAAGGACGGGTCCAGTGCTTCGGCACTGTTGAGTAGAGTGTGAGCTCCGTAACTGGTCGCGTCAC
tggggcacgccgtctggGGGAGACGCGACCGAAATGGTGAAGGACGGGTCCAGTGCTTCGGCACTGTTGAGTAGAGTGTGAGCTCCGTAACTGGTCGCGTCAC
ttactgcgaccgcaataGGGAGACGCGACCGAAATGGTGAAGGACGGGTCCAGTGCTTCGGCACTGTTGAGTAGAGTGTGAGCTCCGTAACTGGTCGCGTCAC
gcgcgcaaccgggtagaGGGAGACGCGACCGAAATGGTGAAGGACGGGTCCAGTGCTTCGGCACTGTTGAGTAGAGTGTGAGCTCCGTAACTGGTCGCGTCAC
gtaactcacggcgctatGGGAGACGCGACCGAAATGGTGAAGGACGGGTCCAGTGCTTCGGCACTGTTGAGTAGAGTGTGAGCTCCGTAACTGGTCGCGTCAC
gctagatagacggcatgGGGAGACGCGACCGAAATGGTGAAGGACGGGTCCAGTGCTTCGGCACTGTTGAGTAGAGTGTGAGCTCCGTAACTGGTCGCGTCAC
gtaagatgctccggttaGGGAGACGCGACCGAAATGGTGAAGGACGGGTCCAGTGCTTCGGCACTGTTGAGTAGAGTGTGAGCTCCGTAACTGGTCGCGTCAC
tgatgtaccgttgagcaGGGAGACGCGACCGAAATGGTGAAGGACGGGTCCAGTGCTTCGGCACTGTTGAGTAGAGTGTGAGCTCCGTAACTGGTCGCGTCAC
tcgctagagcatggtttGGGAGACGCGACCGAAATGGTGAAGGACGGGTCCAGTGCTTCGGCACTGTTGAGTAGAGTGTGAGCTCCGTAACTGGTCGCGTCAC
tggggcacgccgtctggGGGAGACGCGACCGAAATGGTGAAGGACGGGTCCAGTGCTTCGGCACTGTTGAGTAGAGTGTGAGCTCCGTAACTGGTCGCGTCAC
ttactgcgaccgcaataGGGAGACGCGACCGAAATGGTGAAGGACGGGTCCAGTGCTTCGGCACTGTTGAGTAGAGTGTGAGCTCCGTAACTGGTCGCGTCAC
gcgcgcaaccgggtagaGGGAGACGCGACCGAAATGGTGAAGGACGGGTCCAGTGCTTCGGCACTGTTGAGTAGAGTGTGAGCTCCGTAACTGGTCGCGTCAC
gtaactcacggcgctatGGGAGACGCGACCGAAATGGTGAAGGACGGGTCCAGTGCTTCGGCACTGTTGAGTAGAGTGTGAGCTCCGTAACTGGTCGCGTCAC
gctagatagacggcatgGGGAGACGCGACCGAAATGGTGAAGGACGGGTCCAGTGCTTCGGCACTGTTGAGTAGAGTGTGAGCTCCGTAACTGGTCGCGTCAC
gtaagatgctccggttaGGGAGACGCGACCGAAATGGTGAAGGACGGGTCCAGTGCTTCGGCACTGTTGAGTAGAGTGTGAGCTCCGTAACTGGTCGCGTCAC
tgatgtaccgttgagcaGGGAGACGCGACCGAAATGGTGAAGGACGGGTCCAGTGCTTCGGCACTGTTGAGTAGAGTGTGAGCTCCGTAACTGGTCGCGTCAC
tcgctagagcatggtttGGGAGACGCGACCGAAATGGTGAAGGACGGGTCCAGTGCTTCGGCACTGTTGAGTAGAGTGTGAGCTCCGTAACTGGTCGCGTCAC
tggggcacgccgtctggGGGAGACGCGACCGAAATGGTGAAGGACGGGTCCAGTGCTTCGGCACTGTTGAGTAGAGTGTGAGCTCCGTAACTGGTCGCGTCAC
ttactgcgaccgcaataGGGAGACGCGACCGAAATGGTGAAGGACGGGTCCAGTGCTTCGGCACTGTTGAGTAGAGTGTGAGCTCCGTAACTGGTCGCGTCAC
gcgcgcaaccgggtagaGGGAGACGCGACCGAAATGGTGAAGGACGGGTCCAGTGCTTCGGCACTGTTGAGTAGAGTGTGAGCTCCGTAACTGGTCGCGTCAC
gtaactcacggcgctatGGGAGACGCGACCGAAATGGTGAAGGACGGGTCCAGTGCTTCGGCACTGTTGAGTAGAGTGTGAGCTCCGTAACTGGTCGCGTCAC
gctagatagacggcatgGGGAGACGCGACCGAAATGGTGAAGGACGGGTCCAGTGCTTCGGCACTGTTGAGTAGAGTGTGAGCTCCGTAACTGGTCGCGTCAC
gtaagatgctccggttaGGGAGACGCGACCGAAATGGTGAAGGACGGGTCCAGTGCTTCGGCACTGTTGAGTAGAGTGTGAGCTCCGTAACTGGTCGCGTCAC
tgatgtaccgttgagcaGGGAGACGCGACCGAAATGGTGAAGGACGGGTCCAGTGCTTCGGCACTGTTGAGTAGAGTGTGAGCTCCGTAACTGGTCGCGTCAC
tcgctagagcatggtttGGGAGACGCGACCGAAATGGTGAAGGACGGGTCCAGTGCTTCGGCACTGTTGAGTAGAGTGTGAGCTCCGTAACTGGTCGCGTCAC
tggggcacgccgtctggGGGAGACGCGACCGAAATGGTGAAGGACGGGTCCAGTGCTTCGGCACTGTTGAGTAGAGTGTGAGCTCCGTAACTGGTCGCGTCAC
ttactgcgaccgcaataGGGAGACGCGACCGAAATGGTGAAGGACGGGTCCAGTGCTTCGGCACTGTTGAGTAGAGTGTGAGCTCCGTAACTGGTCGCGTCAC
gcgcgcaaccgggtagaGGGAGACGCGACCGAAATGGTGAAGGACGGGTCCAGTGCTTCGGCACTGTTGAGTAGAGTGTGAGCTCCGTAACTGGTCGCGTCAC
gtaactcacggcgctatGGGAGACGCGACCGAAATGGTGAAGGACGGGTCCAGTGCTTCGGCACTGTTGAGTAGAGTGTGAGCTCCGTAACTGGTCGCGTCAC
gctagatagacggcatgGGGAGACGCGACCGAAATGGTGAAGGACGGGTCCAGTGCTTCGGCACTGTTGAGTAGAGTGTGAGCTCCGTAACTGGTCGCGTCAC
gtaagatgctccggttaGGGAGACGCGACCGAAATGGTGAAGGACGGGTCCAGTGCTTCGGCACTGTTGAGTAGAGTGTGAGCTCCGTAACTGGTCGCGTCAC
tgatgtaccgttgagcaGGGAGACGCGACCGAAATGGTGAAGGACGGGTCCAGTGCTTCGGCACTGTTGAGTAGAGTGTGAGCTCCGTAACTGGTCGCGTCAC
tcgctagagcatggtttGGGAGACGCGACCGAAATGGTGAAGGACGGGTCCAGTGCTTCGGCACTGTTGAGTAGAGTGTGAGCTCCGTAACTGGTCGCGTCAC
tggggcacgccgtctggGGGAGACGCGACCGAAATGGTGAAGGACGGGTCCAGTGCTTCGGCACTGTTGAGTAGAGTGTGAGCTCCGTAACTGGTCGCGTCAC
ttactgcgaccgcaataGGGAGACGCGACCGAAATGGTGAAGGACGGGTCCAGTGCTTCGGCACTGTTGAGTAGAGTGTGAGCTCCGTAACTGGTCGCGTCAC
gcgcgcaaccgggtagaGGGAGACGCGACCGAAATGGTGAAGGACGGGTCCAGTGCTTCGGCACTGTTGAGTAGAGTGTGAGCTCCGTAACTGGTCGCGTCAC
gtaactcacggcgctatGGGAGACGCGACCGAAATGGTGAAGGACGGGTCCAGTGCTTCGGCACTGTTGAGTAGAGTGTGAGCTCCGTAACTGGTCGCGTCAC
gctagatagacggcatgGGGAGACGCGACCGAAATGGTGAAGGACGGGTCCAGTGCTTCGGCACTGTTGAGTAGAGTGTGAGCTCCGTAACTGGTCGCGTCAC
gtaagatgctccggttaGGGAGACGCGACCGAAATGGTGAAGGACGGGTCCAGTGCTTCGGCACTGTTGAGTAGAGTGTGAGCTCCGTAACTGGTCGCGTCAC
tgatgtaccgttgagcaGGGAGACGCGACCGAAATGGTGAAGGACGGGTCCAGTGCTTCGGCACTGTTGAGTAGAGTGTGAGCTCCGTAACTGGTCGCGTCAC
tcgctagagcatggtttGGGAGACGCGACCGAAATGGTGAAGGACGGGTCCAGTGCTTCGGCACTGTTGAGTAGAGTGTGAGCTCCGTAACTGGTCGCGTCAC
tggggcacgccgtctggGGGAGACGCGACCGAAATGGTGAAGGACGGGTCCAGTGCTTCGGCACTGTTGAGTAGAGTGTGAGCTCCGTAACTGGTCGCGTCAC
ttactgcgaccgcaataGGGAGACGCGACCGAAATGGTGAAGGACGGGTCCAGTGCTTCGGCACTGTTGAGTAGAGTGTGAGCTCCGTAACTGGTCGCGTCAC
gcgcgcaaccgggtagaGGGAGACGCGACCGAAATGGTGAAGGACGGGTCCAGTGCTTCGGCACTGTTGAGTAGAGTGTGAGCTCCGTAACTGGTCGCGTCAC
gtaactcacggcgctatGGGAGACGCGACCGAAATGGTGAAGGACGGGTCCAGTGCTTCGGCACTGTTGAGTAGAGTGTGAGCTCCGTAACTGGTCGCGTCAC
gctagatagacggcatgGGGAGACGCGACCGAAATGGTGAAGGACGGGTCCAGTGCTTCGGCACTGTTGAGTAGAGTGTGAGCTCCGTAACTGGTCGCGTCAC
gtaagatgctccggttaGGGAGACGCGACCGAAATGGTGAAGGACGGGTCCAGTGCTTCGGCACTGTTGAGTAGAGTGTGAGCTCCGTAACTGGTCGCGTCAC
tgatgtaccgttgagcaGGGAGACGCGACCGAAATGGTGAAGGACGGGTCCAGTGCTTCGGCACTGTTGAGTAGAGTGTGAGCTCCGTAACTGGTCGCGTCAC
tcgctagagcatggtttGGGAGACGCGACCGAAATGGTGAAGGACGGGTCCAGTGCTTCGGCACTGTTGAGTAGAGTGTGAGCTCCGTAACTGGTCGCGTCAC
tggggcacgccgtctggGGGAGACGCGACCGAAATGGTGAAGGACGGGTCCAGTGCTTCGGCACTGTTGAGTAGAGTGTGAGCTCCGTAACTGGTCGCGTCAC
ttactgcgaccgcaataGGGAGACGCGACCGAAATGGTGAAGGACGGGTCCAGTGCTTCGGCACTGTTGAGTAGAGTGTGAGCTCCGTAACTGGTCGCGTCAC
gcgcgcaaccgggtagaGGGAGACGCGACCGAAATGGTGAAGGACGGGTCCAGTGCTTCGGCACTGTTGAGTAGAGTGTGAGCTCCGTAACTGGTCGCGTCAC
gtaactcacggcgctatGGGAGACGCGACCGAAATGGTGAAGGACGGGTCCAGTGCTTCGGCACTGTTGAGTAGAGTGTGAGCTCCGTAACTGGTCGCGTCAC
GCTAGCGTCCGAAGCTT
```

**Table S2 | Number of helicase trajectories at each condition.**

Cell values: $N_{sense}$ / $N_{a\text{-}sense}$

| ATP (μM) | ADP (μM) | Force-Oppose Translocase | | | | Force-Assist Helicase | | | | Force-Assist Translocase | | | |
|---|---|---|---|---|---|---|---|---|---|---|---|---|---|
| | Applied force (pN) | 36 | 28 | 20 | 12 | 12 | 20 | 28 | 36 | 12 | 20 | 28 | 36 |
| 3000 | 0 | 12 / 0 | 10 / 0 | 9 / 0 | 10 / 0 | 2 / 32 | 2 / 31 | 2 / 33 | 2 / 38 | 13 / 0 | 13 / 0 | 14 / 0 | 13 / 0 |
| 1000 | 0 | 89 / 0 | 79 / 0 | 74 / 0 | 70 / 0 | 35 / 0 | 39 / 0 | 44 / 0 | 60 / 0 | 0 / 10 | 0 / 13 | 0 / 13 | 0 / 13 |
| 100 | 0 | 89 / 0 | 80 / 0 | 68 / 0 | 58 / 0 | 11 / 1 | 18 / 0 | 24 / 0 | 34 / 4 | 4 / 1 | 4 / 4 | 4 / 3 | 5 / 6 |
| 10 | 0 | 24 / 0 | 21 / 0 | 22 / 0 | 22 / 0 | 9 / 2 | 13 / 1 | 16 / 1 | 33 / 14 | 6 / 2 | 6 / 1 | 6 / 2 | 7 / 4 |
| 5 | 0 | 12 / 0 | 12 / 0 | 12 / 0 | 10 / 0 | 31 / 0 | 37 / 0 | 42 / 0 | 45 / 0 | 0 / 8 | 0 / 10 | 0 / 11 | 0 / 15 |
| 1 | 0 | 27 / 0 | 23 / 0 | 22 / 0 | 21 / 0 | 18 / 0 | 24 / 0 | 27 / 5 | 37 / 22 | 5 / 0 | 5 / 2 | 6 / 3 | 6 / 3 |
| 0 | 0 | 0 / 0 | 0 / 0 | 0 / 0 | 0 / 0 | 3 / 0 | 4 / 0 | 6 / 0 | 10 / 0 | 0 / 0 | 0 / 0 | 0 / 0 | 0 / 0 |
| 10 | 0 | 24 / 0 | 21 / 0 | 22 / 0 | 22 / 0 | 9 / 2 | 13 / 1 | 16 / 1 | 33 / 14 | 6 / 2 | 6 / 1 | 6 / 2 | 7 / 4 |
| 10 | 10 | 49 / 0 | 37 / 0 | 29 / 0 | 28 / 0 | 20 / 0 | 22 / 0 | 28 / 0 | 40 / 0 | 0 / 8 | 0 / 10 | 0 / 10 | 0 / 8 |
| 10 | 50 | 29 / 0 | 29 / 0 | 30 / 0 | 29 / 0 | 38 / 5 | 44 / 5 | 55 / 6 | 67 / 33 | 8 / 6 | 8 / 6 | 8 / 5 | 8 / 7 |
| 10 | 250 | 18 / 0 | 18 / 0 | 16 / 0 | 15 / 0 | 12 / 1 | 14 / 1 | 15 / 1 | 20 / 3 | 0 / 0 | 0 / 2 | 0 / 3 | 0 / 0 |
| 10 | 1000 | 4 / 0 | 4 / 0 | 4 / 0 | 4 / 0 | 9 / 0 | 11 / 0 | 11 / 0 | 11 / 0 | 0 / 2 | 0 / 2 | 0 / 2 | 0 / 1 |

**Table S2:** This table shows the number of individual helicase trajectories for each condition on each strand of the DNA test track (sense/antisense). Note the condition [ATP] = 10 μM, [ADP] = 0 is duplicated within the table for ease of comparison in the ATP titration experiment and ADP titration experiment.

**Table S3 | Number of total steps observed at each condition.**

Cell values: $N_{sense}$ / $N_{a\text{-}sense}$

| ATP (μM) | ADP (μM) | Force-Oppose Translocase | | | | Force-Assist Helicase | | | | Force-Assist Translocase | | | |
|---|---|---|---|---|---|---|---|---|---|---|---|---|---|
| | Applied force (pN) | 36 | 28 | 20 | 12 | 12 | 20 | 28 | 36 | 12 | 20 | 28 | 36 |
| 3000 | 0 | 4727 / 0 | 13602 / 0 | 17491 / 0 | 30760 / 0 | 2163 / 27710 | 2146 / 24784 | 1280 / 25216 | 656 / 42803 | 12071 / 0 | 8623 / 0 | 10608 / 0 | 12172 / 0 |
| 1000 | 0 | 77410 / 0 | 85651 / 0 | 75201 / 0 | 76023 / 0 | 27841 / 0 | 36747 / 0 | 40495 / 0 | 62828 / 0 | 0 / 10260 | 0 / 24589 | 0 / 16560 | 0 / 16916 |
| 100 | 0 | 73587 / 0 | 81886 / 0 | 56366 / 0 | 76044 / 0 | 9996 / 61 | 13791 / 0 | 19809 / 0 | 27316 / 5143 | 4080 / 1021 | 3467 / 12007 | 3991 / 4635 | 4569 / 15710 |
| 10 | 0 | 8808 / 0 | 24181 / 0 | 21146 / 0 | 26880 / 0 | 9488 / 3795 | 13592 / 1587 | 15677 / 824 | 35760 / 26234 | 8276 / 2162 | 6775 / 913 | 7484 / 4306 | 8831 / 4802 |
| 5 | 0 | 5208 / 0 | 13645 / 0 | 13223 / 0 | 15142 / 0 | 21812 / 0 | 29749 / 0 | 27778 / 0 | 25236 / 0 | 0 / 9863 | 0 / 7760 | 0 / 10897 | 0 / 22106 |
| 1 | 0 | 2979 / 0 | 8862 / 0 | 15746 / 0 | 17586 / 0 | 4164 / 0 | 11069 / 0 | 17254 / 3859 | 18559 / 32980 | 5403 / 0 | 3908 / 1303 | 4295 / 5709 | 4458 / 510 |
| 0 | 0 | 0 / 0 | 0 / 0 | 0 / 0 | 0 / 0 | 158 / 0 | 667 / 0 | 2064 / 0 | 2613 / 0 | 0 / 0 | 0 / 0 | 0 / 0 | 0 / 0 |
| 10 | 0 | 8808 / 0 | 24181 / 0 | 21146 / 0 | 26880 / 0 | 9488 / 3795 | 13592 / 1587 | 15677 / 824 | 35760 / 26234 | 8276 / 2162 | 6775 / 913 | 7484 / 4306 | 8831 / 4802 |
| 10 | 10 | 30981 / 0 | 38790 / 0 | 46380 / 0 | 38217 / 0 | 11330 / 0 | 15993 / 0 | 22239 / 0 | 37109 / 0 | 0 / 7616 | 0 / 10547 | 0 / 13264 | 0 / 13658 |
| 10 | 50 | 11619 / 0 | 27096 / 0 | 26531 / 0 | 29721 / 0 | 16586 / 1181 | 33776 / 1236 | 41585 / 1842 | 45877 / 63897 | 8720 / 5394 | 5983 / 7870 | 8671 / 5809 | 8325 / 5672 |
| 10 | 250 | 5280 / 0 | 14084 / 0 | 13587 / 0 | 11306 / 0 | 9154 / 340 | 12518 / 310 | 11494 / 560 | 12335 / 2607 | 0 / 0 | 0 / 1664 | 0 / 1783 | 0 / 0 |
| 10 | 1000 | 879 / 0 | 679 / 0 | 960 / 0 | 703 / 0 | 5630 / 0 | 5534 / 0 | 6088 / 0 | 5582 / 0 | 0 / 2264 | 0 / 1431 | 0 / 699 | 0 / 1256 |

**Table S3:** Number of total steps observed at each condition on each strand (sense/antisense) including all forwards and backwards steps.

**Construction of 3′ feeding ion-current-to-DNA sequence map.**

The orientation in which DNA is fed into the pore strongly affects the ion currents that are measured during a typical SPRNT experiment[2]. Therefore, there are two separate mappings of ion-current to DNA sequence depending on whether the SPRNT experiment is in a force-assisting (3′ feeding) or force-opposing (5′ feeding) configuration. We have shown previously that predominantly 4 nucleotides centered within the nanopore constriction contribute to the measured ion current. This means that one can map out the currents that result from all 4-nucleotide combinations, dubbed a 'quadromer,' within the pore[3]. The current-to-sequence map for 5′ feeding was previously determined[3] however a complete current-to-sequence map for 3′ feeding was needed to interpret 3′ feeding SPRNT data. We constructed the 3′ feeding current-to-sequence map in a similar manner by measuring each 4-nucleotide combination in a number of different sequence contexts within the phiX174 genome, the results of which are displayed in Table S4. PhiX174 DNA was prepared as previously described in Noakes *et al.*[4]. In nanopore sequencing, these ion-current-to-DNA-sequence mappings are used to sequence the DNA. In SPRNT, these mappings can be used to predict the ion current that will be measured for a given DNA sequence and to align the measured ion-currents to the known DNA sequence.

**Construction of ion current consensuses and alignment to DNA sequence within MspA.**

In practice, there are slight deviations of the measured ion current from that which would be predicted from the known sequence. This is because other bases outside of the quadromer contribute to the ion current, albeit to a significantly smaller degree. For the highest resolution in SPRNT, it is best to measure the DNA substrate directly and form a consensus of measured ion-currents to which individual helicase traces can be aligned.

The applied voltage also has a significant effect on the ion currents measured during a SPRNT experiment. As the applied voltage is reduced from 180 mV to 60 mV, there are two principle effects to the ion current: 1) The ion current is reduced due to the reduced potential across the pore and 2) the reduction in applied force relaxes the tension on the DNA between where it is held by the enzyme and where it is pulled on by the electric field within the pore constriction. This second effect reduces the amount by which the DNA is stretched, repositioning the DNA within the pore, thereby changing the position of the DNA as measured by SPRNT[4–6]. At lower voltages (lower applied force) the signal-to-noise ratio is also reduced, limiting the ability to resolve ion current steps.

For these reasons, different ion-current consensuses were measured for each DNA sequence (sense, antisense), each DNA feeding orientation (3′ and 5′ feeding), and each applied voltage (60mV, 100mV, 140mV, and 180mV). Below in Figs. S3-S5, we show the ion-current consensuses for sense and antisense strands for each of the configurations used within this study. The ion currents for the force-opposing antisense strand had a significant number of adjacent indistinguishable ion current levels. This makes it difficult to distinguish enzyme steps for large patches of the antisense sequence making it not well-suited for SPRNT experiments. We have not used any of this data in our analysis. The ion currents are aligned to the DNA sequence within the nanopore constriction responsible for generating those ion currents via comparison to the ion current prediction (red current levels in 180mV data Figs. S3-S5). Gaps within the ion-current traces represent missing ion-current levels, i.e. the number of observed levels is fewer than the known number of DNA bases. The interpretation here is that in regions containing gaps, SPRNT lacks the resolution to resolve individual nucleotide steps, however, the overall pattern of ion-currents still provides the relative DNA position. Regions containing gaps were not used in analysis of single-nucleotide step dwell-times or backstepping.

Raw ion current reads are sampled at 50 kHz and downsampled to 5 kHz for analysis. Data processing is as previously described[3,5,7–9]. Briefly, events containing enzyme-controlled ion-current traces are automatically detected and flagged for analysis with custom software built in Matlab (Mathworks). Events were categorized and sorted by strand and feeding orientation by eye via recognition of distinct ion-current patterns for each strand and DNA orientation (see consensuses below). Individual enzyme traces were then aligned to the ion-current consensuses using a dynamic-programming based alignment code similar to those previously described[3,5] (code included in supplementary code package). Alignments were made to the consensus with gaps removed and then measured positions were corrected by reinserting the gaps to reflect the known DNA sequence alignment. Alignment step parameters (step, backstep, skip, hold, and bad-level) were tuned for each applied force and [ATP] and [ADP] condition such that the proper global alignment was achieved. Alignments were verified by hand for each condition.

**Table S4 | Measured Ion-current values for 3′ feeding.**

| | | | | | | | |
|---|---|---|---|---|---|---|---|
| AAAA | 0.555 ± 0.049 | CAAA | 0.397 ± 0.077 | GAAA | 0.502 ± 0.061 | TAAA | 0.545 ± 0.042 |
| AAAC | 0.484 ± 0.052 | CAAC | 0.342 ± 0.014 | GAAC | 0.417 ± 0.051 | TAAC | 0.506 ± 0.025 |
| AAAG | 0.574 ± 0.054 | CAAG | 0.492 ± 0.079 | GAAG | 0.679 ± 0.039 | TAAG | 0.622 ± 0.036 |
| AAAT | 0.59 ± 0.049 | CAAT | 0.356 ± 0.057 | GAAT | 0.557 ± 0.03 | TAAT | 0.404 ± 0.086 |
| AACA | 0.38 ± 0.041 | CACA | 0.322 ± 0.01 | GACA | 0.456 ± 0.058 | TACA | 0.454 ± 0.028 |
| AACC | 0.318 ± 0.033 | CACC | 0.32 ± 0.066 | GACC | 0.331 ± 0.028 | TACC | 0.313 ± 0.033 |
| AACG | 0.391 ± 0.048 | CACG | 0.3 ± 0.011 | GACG | 0.396 ± 0.047 | TACG | 0.365 ± 0.051 |
| AACT | 0.248 ± 0.024 | CACT | 0.245 ± 0.024 | GACT | 0.278 ± 0.045 | TACT | 0.265 ± 0.026 |
| AAGA | 0.619 ± 0.058 | CAGA | 0.336 ± 0.055 | GAGA | 0.585 ± 0.04 | TAGA | 0.558 ± 0.028 |
| AAGC | 0.523 ± 0.11 | CAGC | 0.432 ± 0.033 | GAGC | 0.463 ± 0.099 | TAGC | 0.403 ± 0.01 |
| AAGG | 0.63 ± 0.061 | CAGG | 0.557 ± 0.045 | GAGG | 0.629 ± 0.059 | TAGG | 0.606 ± 0.022 |
| AAGT | 0.546 ± 0.067 | CAGT | 0.464 ± 0.05 | GAGT | 0.604 ± 0.054 | TAGT | 0.506 ± 0.024 |
| AATA | 0.484 ± 0.1 | CATA | 0.402 ± 0.098 | GATA | 0.552 ± 0.046 | TATA | 0.472 ± 0.05 |
| AATC | 0.471 ± 0.059 | CATC | 0.414 ± 0.074 | GATC | 0.43 ± 0.01 | TATC | 0.469 ± 0.019 |
| AATG | 0.633 ± 0.059 | CATG | 0.48 ± 0.064 | GATG | 0.623 ± 0.051 | TATG | 0.537 ± 0.063 |
| AATT | 0.535 ± 0.043 | CATT | 0.383 ± 0.064 | GATT | 0.522 ± 0.042 | TATT | 0.428 ± 0.063 |
| ACAA | 0.346 ± 0.016 | CCAA | 0.318 ± 0.051 | GCAA | 0.363 ± 0.018 | TCAA | 0.343 ± 0.05 |
| ACAC | 0.318 ± 0.062 | CCAC | 0.274 ± 0.01 | GCAC | 0.303 ± 0.034 | TCAC | 0.295 ± 0.019 |
| ACAG | 0.339 ± 0.016 | CCAG | 0.314 ± 0.018 | GCAG | 0.367 ± 0.026 | TCAG | 0.351 ± 0.041 |
| ACAT | 0.321 ± 0.01 | CCAT | 0.293 ± 0.029 | GCAT | 0.362 ± 0.01 | TCAT | 0.282 ± 0.04 |
| ACCA | 0.329 ± 0.029 | CCCA | 0.277 ± 0.021 | GCCA | 0.293 ± 0.012 | TCCA | 0.287 ± 0.034 |
| ACCC | 0.284 ± 0.024 | CCCC | 0.266 ± 0.015 | GCCC | 0.268 ± 0.01 | TCCC | 0.252 ± 0.015 |
| ACCG | 0.296 ± 0.01 | CCCG | 0.301 ± 0.016 | GCCG | 0.287 ± 0.027 | TCCG | 0.296 ± 0.013 |
| ACCT | 0.288 ± 0.04 | CCCT | 0.249 ± 0.015 | GCCT | 0.3 ± 0.01 | TCCT | 0.254 ± 0.026 |
| ACGA | 0.352 ± 0.012 | CCGA | 0.335 ± 0.021 | GCGA | 0.348 ± 0.025 | TCGA | 0.335 ± 0.01 |
| ACGC | 0.347 ± 0.029 | CCGC | 0.361 ± 0.066 | GCGC | 0.265 ± 0.034 | TCGC | 0.31 ± 0.018 |
| ACGG | 0.345 ± 0.022 | CCGG | 0.282 ± 0.01 | GCGG | 0.318 ± 0.063 | TCGG | 0.34 ± 0.01 |
| ACGT | 0.322 ± 0.035 | CCGT | 0.32 ± 0.035 | GCGT | 0.327 ± 0.026 | TCGT | 0.316 ± 0.012 |
| ACTA | 0.244 ± 0.02 | CCTA | 0.262 ± 0.016 | GCTA | 0.225 ± 0.07 | TCTA | 0.229 ± 0.013 |
| ACTC | 0.227 ± 0.011 | CCTC | 0.236 ± 0.02 | GCTC | 0.213 ± 0.013 | TCTC | 0.231 ± 0.01 |
| ACTG | 0.24 ± 0.016 | CCTG | 0.27 ± 0.16 | GCTG | 0.255 ± 0.063 | TCTG | 0.239 ± 0.015 |
| ACTT | 0.236 ± 0.017 | CCTT | 0.262 ± 0.024 | GCTT | 0.224 ± 0.029 | TCTT | 0.232 ± 0.013 |
| AGAA | 0.541 ± 0.03 | CGAA | 0.604 ± 0.029 | GGAA | 0.547 ± 0.033 | TGAA | 0.52 ± 0.046 |
| AGAC | 0.483 ± 0.061 | CGAC | 0.32 ± 0.025 | GGAC | 0.346 ± 0.055 | TGAC | 0.371 ± 0.045 |
| AGAG | 0.623 ± 0.037 | CGAG | 0.501 ± 0.076 | GGAG | 0.517 ± 0.071 | TGAG | 0.534 ± 0.082 |
| AGAT | 0.513 ± 0.055 | CGAT | 0.493 ± 0.037 | GGAT | 0.522 ± 0.019 | TGAT | 0.568 ± 0.057 |
| AGCA | 0.4 ± 0.034 | CGCA | 0.31 ± 0.04 | GGCA | 0.321 ± 0.042 | TGCA | 0.358 ± 0.016 |
| AGCC | 0.382 ± 0.024 | CGCC | 0.301 ± 0.035 | GGCC | 0.326 ± 0.014 | TGCC | 0.293 ± 0.023 |
| AGCG | 0.369 ± 0.023 | CGCG | 0.33 ± 0.012 | GGCG | 0.314 ± 0.028 | TGCG | 0.32 ± 0.04 |
| AGCT | 0.259 ± 0.038 | CGCT | 0.246 ± 0.023 | GGCT | 0.239 ± 0.027 | TGCT | 0.237 ± 0.033 |
| AGGA | 0.534 ± 0.081 | CGGA | 0.443 ± 0.022 | GGGA | 0.549 ± 0.18 | TGGA | 0.441 ± 0.048 |
| AGGC | 0.383 ± 0.12 | CGGC | 0.35 ± 0.01 | GGGC | 0.4 ± 0.016 | TGGC | 0.316 ± 0.065 |
| AGGG | 0.56 ± 0.069 | CGGG | 0.323 ± 0.01 | GGGG | 0.561 ± 0.047 | TGGG | 0.541 ± 0.073 |
| AGGT | 0.526 ± 0.023 | CGGT | 0.412 ± 0.067 | GGGT | 0.451 ± 0.024 | TGGT | 0.471 ± 0.045 |
| AGTA | 0.54 ± 0.045 | CGTA | 0.457 ± 0.061 | GGTA | 0.504 ± 0.053 | TGTA | 0.572 ± 0.043 |
| AGTC | 0.465 ± 0.096 | CGTC | 0.345 ± 0.039 | GGTC | 0.408 ± 0.058 | TGTC | 0.364 ± 0.046 |
| AGTG | 0.56 ± 0.045 | CGTG | 0.319 ± 0.01 | GGTG | 0.501 ± 0.058 | TGTG | 0.513 ± 0.062 |
| AGTT | 0.463 ± 0.034 | CGTT | 0.409 ± 0.031 | GGTT | 0.441 ± 0.029 | TGTT | 0.492 ± 0.06 |
| ATAA | 0.587 ± 0.036 | CTAA | 0.431 ± 0.098 | GTAA | 0.531 ± 0.01 | TTAA | 0.531 ± 0.084 |
| ATAC | 0.496 ± 0.043 | CTAC | 0.368 ± 0.087 | GTAC | 0.443 ± 0.058 | TTAC | 0.341 ± 0.042 |
| ATAG | 0.55 ± 0.01 | CTAG | 0.215 ± 0.018 | GTAG | 0.55 ± 0.036 | TTAG | 0.595 ± 0.014 |
| ATAT | 0.493 ± 0.059 | CTAT | 0.295 ± 0.069 | GTAT | 0.493 ± 0.05 | TTAT | 0.458 ± 0.059 |
| ATCA | 0.379 ± 0.023 | CTCA | 0.22 ± 0.055 | GTCA | 0.317 ± 0.028 | TTCA | 0.351 ± 0.041 |
| ATCC | 0.291 ± 0.068 | CTCC | 0.27 ± 0.018 | GTCC | 0.307 ± 0.019 | TTCC | 0.311 ± 0.025 |
| ATCG | 0.359 ± 0.02 | CTCG | 0.319 ± 0.026 | GTCG | 0.339 ± 0.014 | TTCG | 0.356 ± 0.022 |
| ATCT | 0.3 ± 0.033 | CTCT | 0.253 ± 0.092 | GTCT | 0.245 ± 0.03 | TTCT | 0.277 ± 0.024 |
| ATGA | 0.652 ± 0.042 | CTGA | 0.392 ± 0.069 | GTGA | 0.496 ± 0.03 | TTGA | 0.5 ± 0.067 |
| ATGC | 0.36 ± 0.049 | CTGC | 0.266 ± 0.022 | GTGC | 0.298 ± 0.036 | TTGC | 0.391 ± 0.064 |
| ATGG | 0.603 ± 0.063 | CTGG | 0.404 ± 0.063 | GTGG | 0.377 ± 0.047 | TTGG | 0.511 ± 0.048 |
| ATGT | 0.561 ± 0.065 | CTGT | 0.471 ± 0.091 | GTGT | 0.423 ± 0.086 | TTGT | 0.483 ± 0.049 |
| ATTA | 0.498 ± 0.035 | CTTA | 0.249 ± 0.076 | GTTA | 0.473 ± 0.06 | TTTA | 0.405 ± 0.034 |
| ATTC | 0.385 ± 0.055 | CTTC | 0.299 ± 0.035 | GTTC | 0.387 ± 0.049 | TTTC | 0.291 ± 0.028 |
| ATTG | 0.506 ± 0.022 | CTTG | 0.366 ± 0.053 | GTTG | 0.484 ± 0.046 | TTTG | 0.41 ± 0.043 |
| ATTT | 0.466 ± 0.048 | CTTT | 0.318 ± 0.052 | GTTT | 0.416 ± 0.046 | TTTT | 0.393 ± 0.033 |

Each quadromer and its measured ion-current value in normalized ion current. Quadromer sequences are written 5′ to 3′.

### Figure S3 | Ion Current Consensus 5′ feeding Sense Strand.

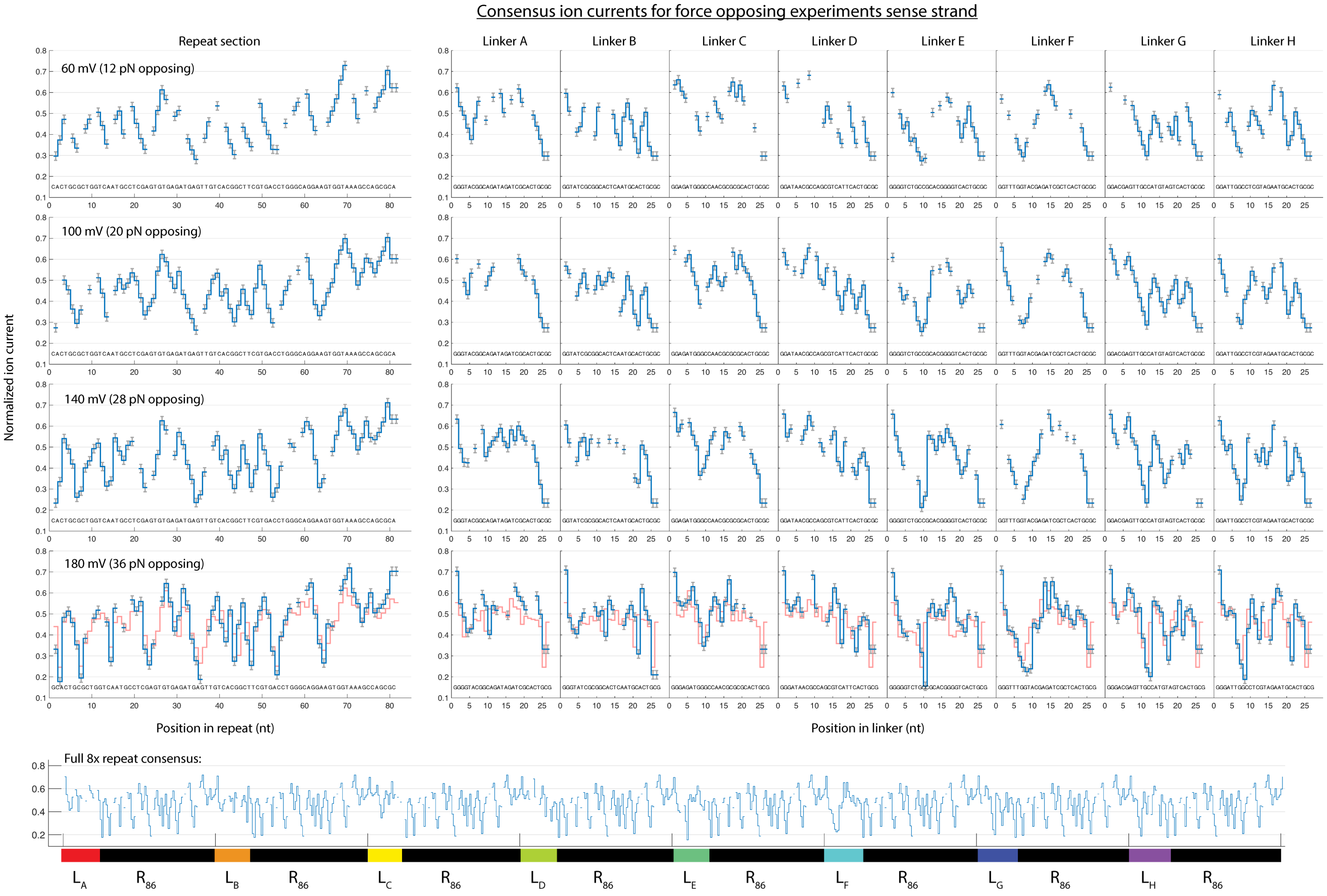

* Ion current levels and DNA sequence shown in the order of appearance in SPRNT traces 3′ to 5′ since PcrA walks from 3′ to 5′ along DNA.

### Figure S4 | Ion Current Consensus 3′ feeding Sense Strand.

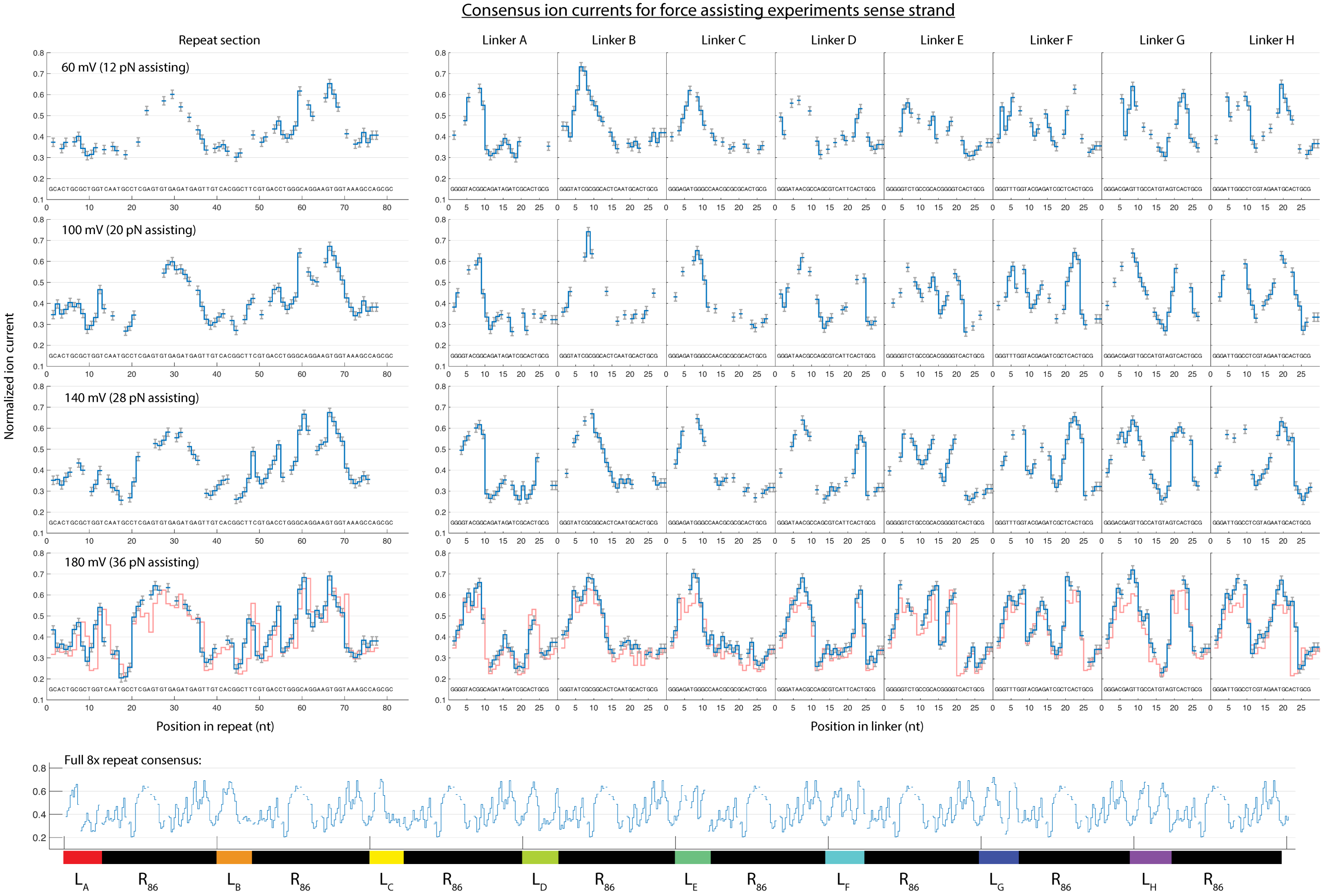

* Ion current levels and DNA sequence shown in the order of appearance in SPRNT traces 3′ to 5′ since PcrA walks from 3′ to 5′ along DNA.

### Figure S5 | Ion Current Consensus 3′ feeding Antisense Strand.

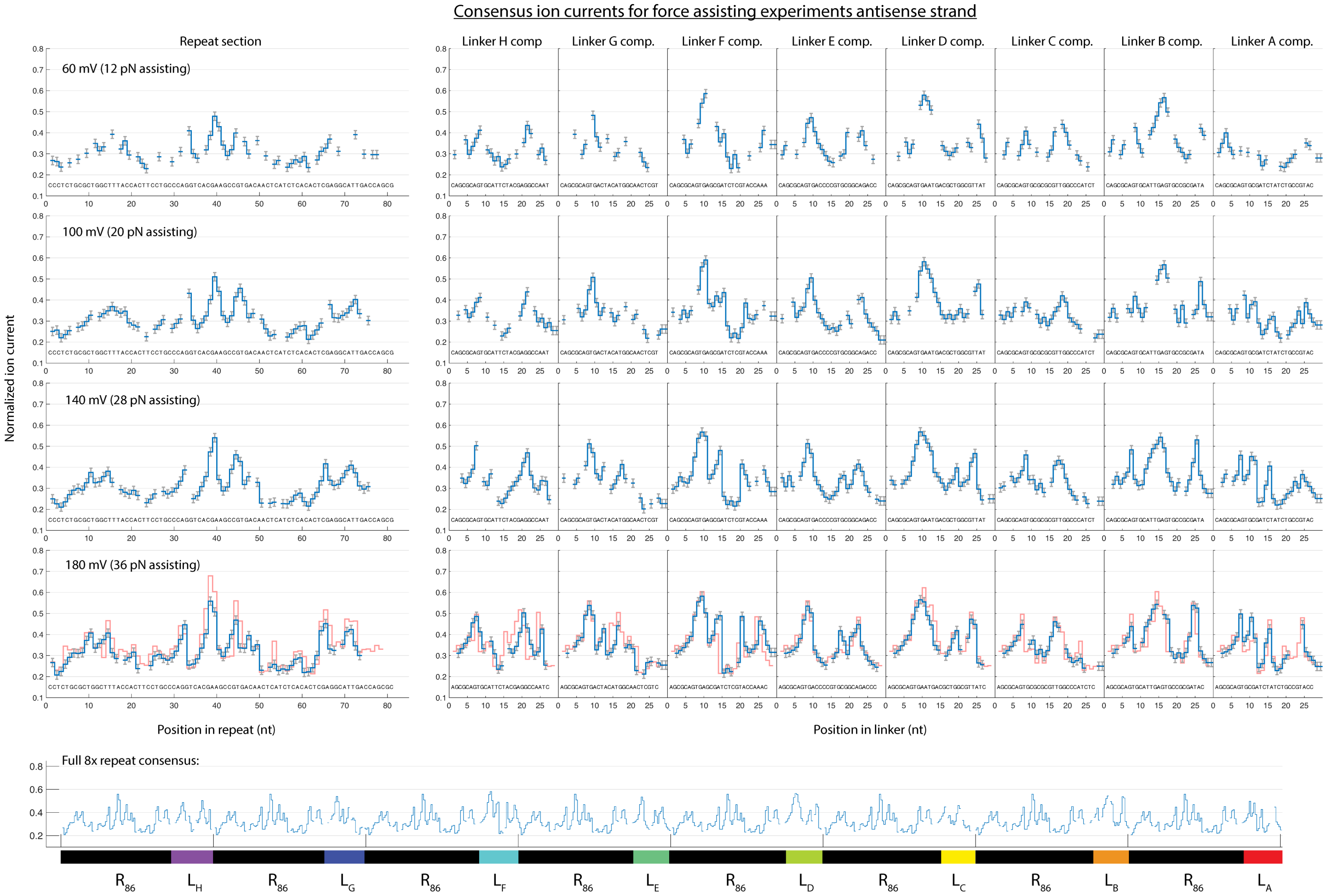

* Ion current levels and DNA sequence shown in the order of appearance in SPRNT traces 3′ to 5′ since PcrA walks from 3′ to 5′ along DNA.

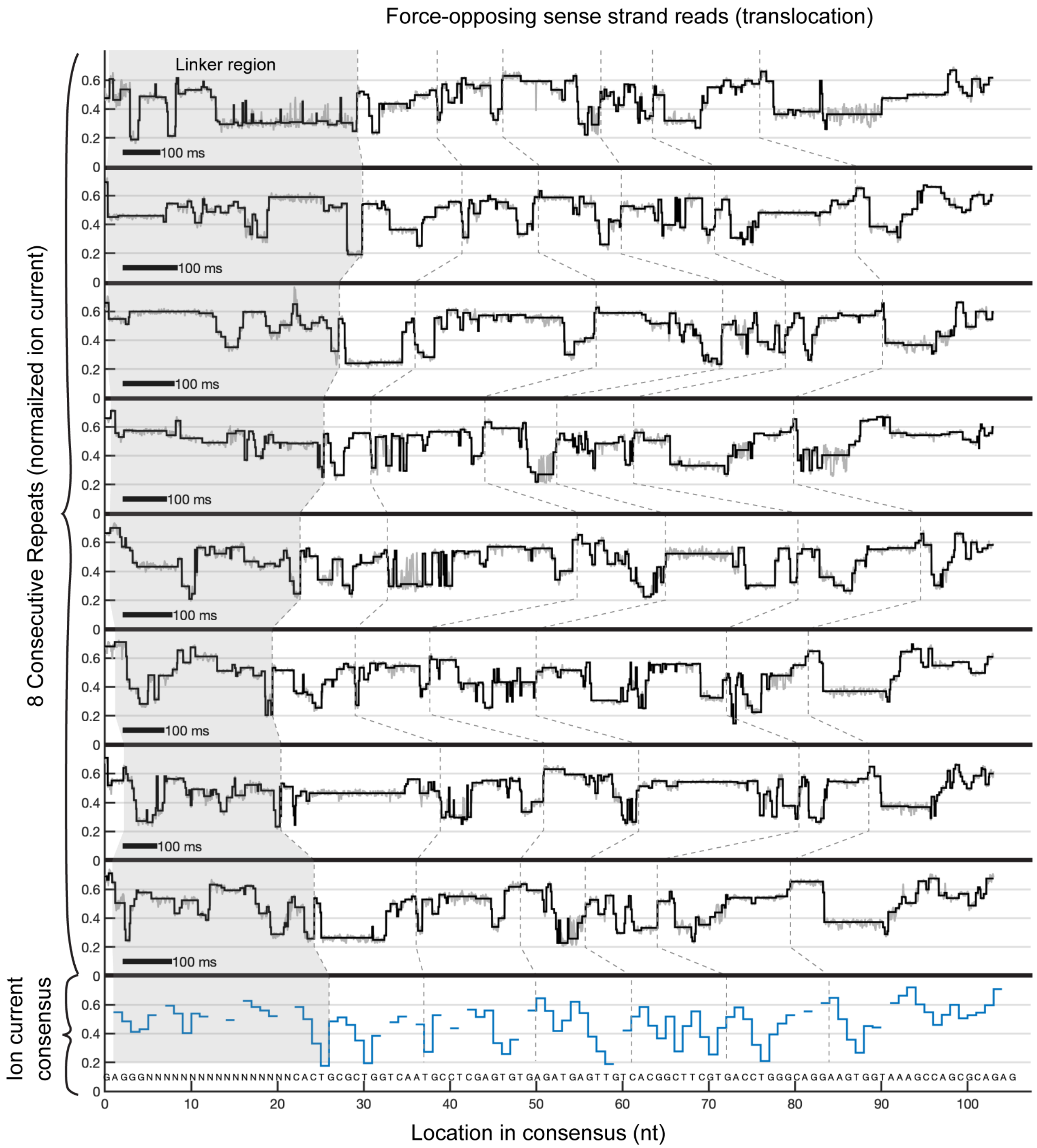

**Figure S6 | Example data traces for Force-Opposing Translocation.**
Eight consecutive repeats within an example data trace with 180mV applied (36pN) downsampled to 500 Hz. The bottom graph shows the empirically determined ion-current consensus for the sense strand 5' feeding with 180 mV applied. Gaps within the ion current consensus represent missing ion current levels that are not resolved because adjacent levels have ion-currents that are too close to be resolved. Each trace begins with a linker region (shaded in gray) that is unique and different for all 8 segments and then continues on through the 86 nt repeat DNA sequence. Dashed lines are guides to the eye marking repeated features within each data trace and the corresponding ion current levels within the ion current consensus. Levels within the ion current trace are automatically detected by a change-point algorithm previously described[6]. Extracted levels are then aligned automatically to the ion current consensus using a dynamic programming algorithm[6]. This representation makes it clear that particular locations tend to have longer dwell-times on average than other locations. Similarly, some locations are more prone to backstepping.

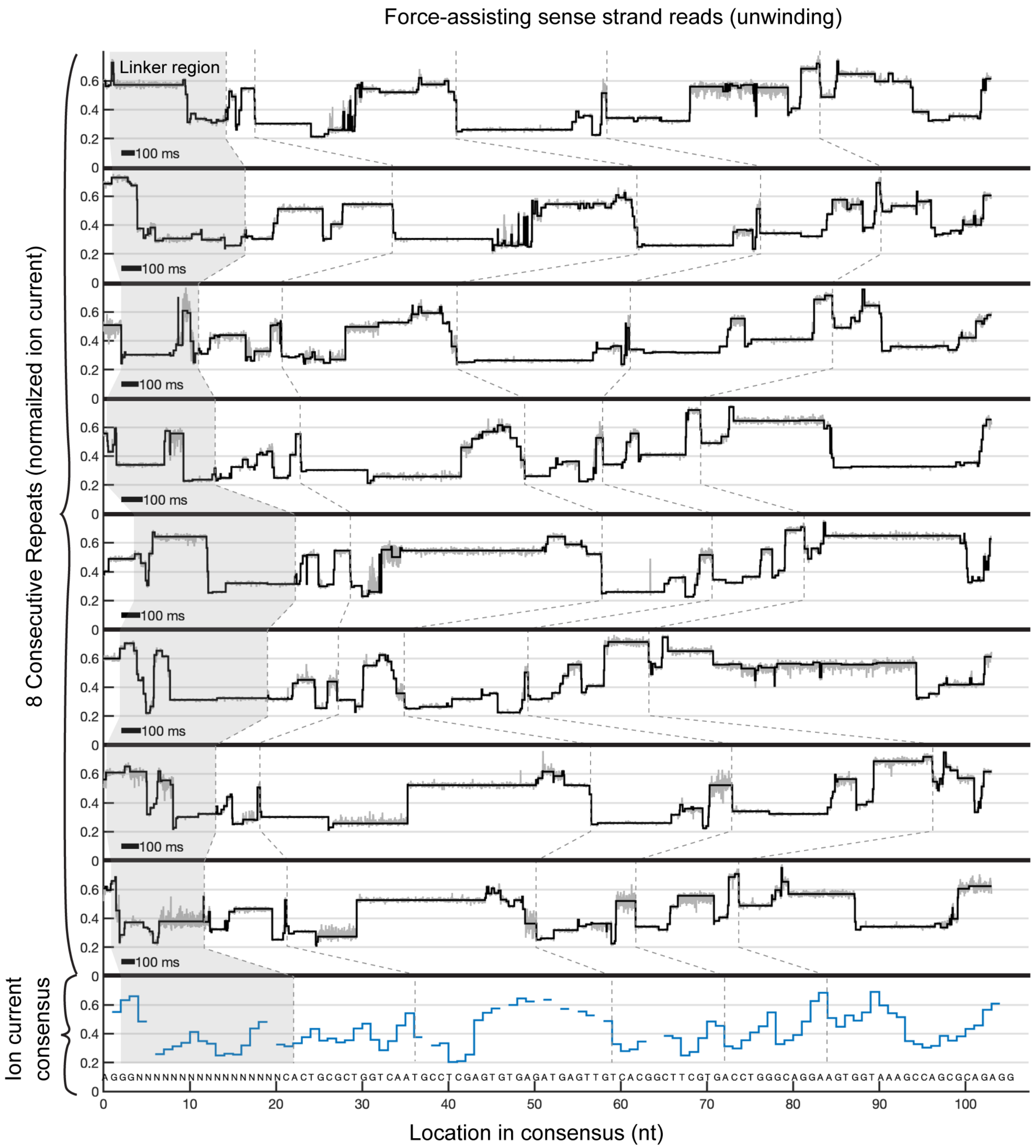

**Figure S7 | Example data traces for Force-Assisting Unwinding.**
Eight consecutive repeats within an example data trace with 180mV applied (36pN) downsampled to 500 Hz. The bottom graph shows the empirically determined ion-current consensus for the sense strand 3' feeding with 180 mV applied. Gaps within the ion current consensus represent missing ion current levels that are not resolved because adjacent levels have ion-currents that are too close to be resolved. Each trace begins with a linker region (shaded in gray) that is unique and different for all 8 segments and then continues on through the 86 nt repeat DNA sequence. Dashed lines are guides to the eye marking repeated features within each data trace and the corresponding ion current levels within the ion current consensus. Levels within the ion current trace are automatically detected by a change-point algorithm previously described[6]. Extracted levels are then aligned automatically to the ion current consensus using a dynamic programming algorithm[6]. This representation makes it clear that particular locations tend to have longer dwell-times on average than other locations. Similarly, some locations are more prone to backstepping.

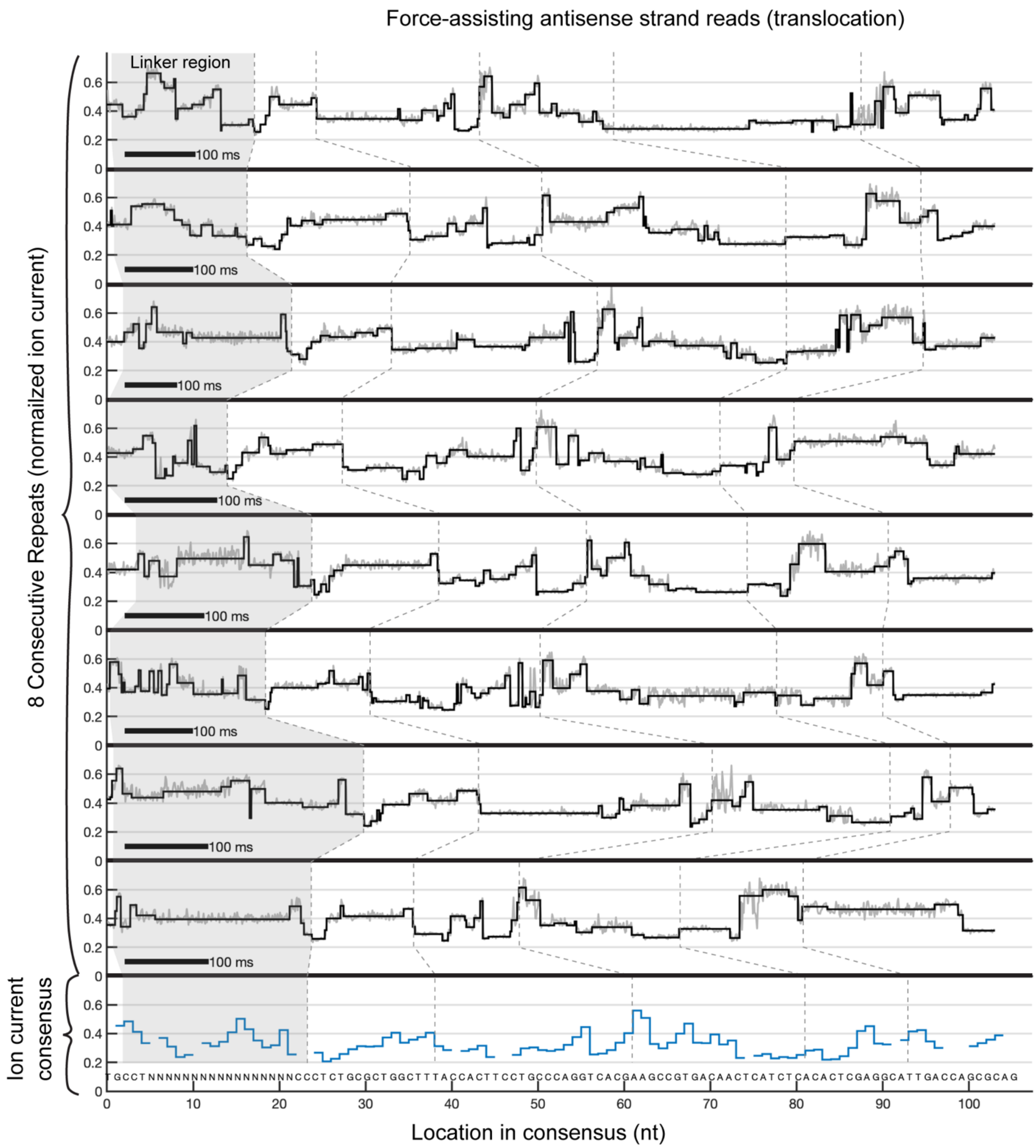

**Figure S8 | Example data traces for Force-Assisting Translocation.**
Eight consecutive repeats within an example data trace with 180mV applied (36pN) downsampled to 500 Hz. The bottom graph shows the empirically determined ion-current consensus for the antisense strand, 3’ feeding with 180 mV applied. Gaps within the ion current consensus represent missing ion current levels that are not resolved because adjacent levels have ion-currents that are too close to be resolved. Each trace begins with a linker region (shaded in gray) that is unique and different for all 8 segments and then continues on through the 86 nt repeat DNA sequence. Dashed lines are guides to the eye marking repeated features within each data trace and the corresponding ion current levels within the ion current consensus. Levels within the ion current trace are automatically detected by a change-point algorithm previously described[6]. Extracted levels are then aligned automatically to the ion current consensus using a dynamic programming algorithm[6]. This representation makes it clear that particular locations tend to have longer dwell-times on average than other locations. Similarly, some locations are more prone to backstepping.

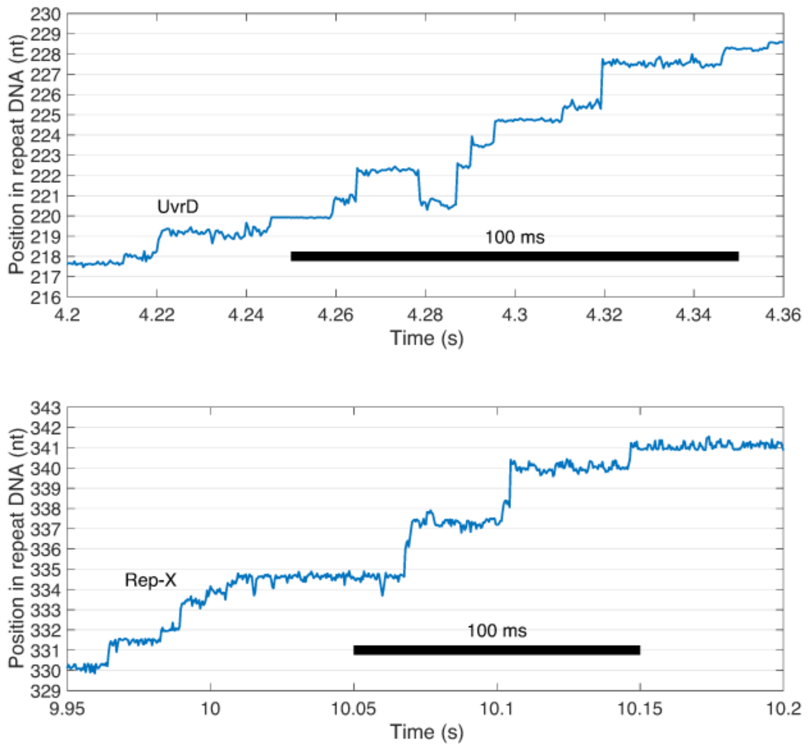

**Figure S9 | Single-nucleotide steps of other SF1 helicases.** Two other SF1 helicases have been run on the SPRNT setup with the repetitive DNA sequence described here: Rep-X helicase[10] at 17°C and UvrD helicase (Quidel Corporation, San Diego) at room temperature and 30mM KCl. Both datasets show single-nucleotide steps at roughly 100 nt/s. This is in contrast to Hel308 helicase, from SF2, which feeds DNA through the pore in two sub-steps per nucleotide. We suspect that the structural similarity of PcrA, Rep, and UvrD causes the three helicases to rest on the pore rim in a similar way and result in only one observable step per nucleotide in contrast to the two steps per nucleotide observed with SF2 helicase Hel308.

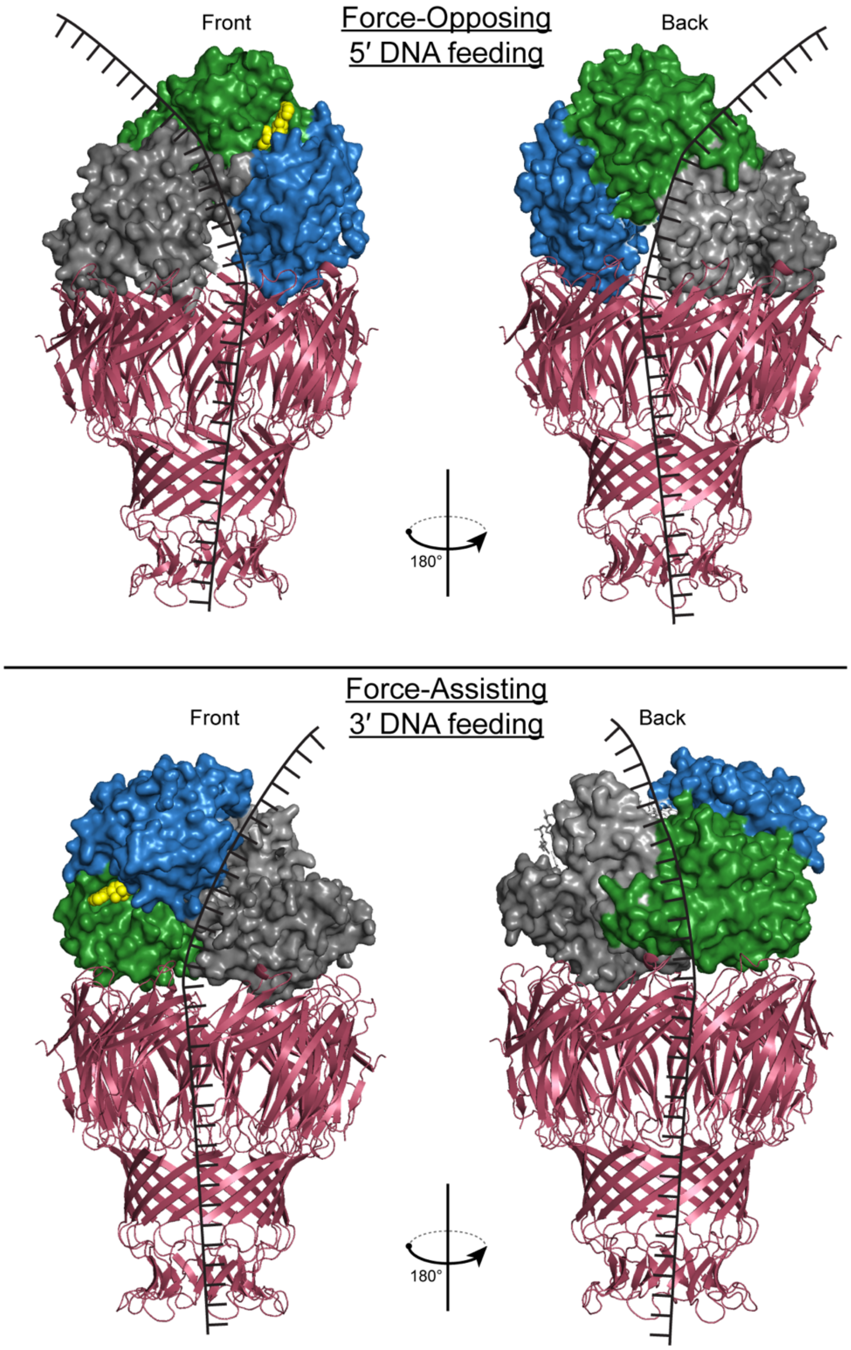

**Figure S10 | Schematic representation of PcrA resting on MspA.** This mock-up is based on crystal structures of PcrA (3PJR) and MspA (1UUN) and represents a best guess for how PcrA rests on top of the MspA pore rim during SPRNT experiments for opposing forces (top) and assisting forces (bottom). Of note is that in each configuration only one of the RecA-like domains 1A (green) or 2A (blue) rests on the pore rim. SPRNT measures the motion of DNA relative to the MspA constriction. Assuming the surface in contact with the pore rim is unchanged as the ATP hydrolysis cycle progresses, we expect that SPRNT measures, in effect, motions of DNA relative to domains in contact with MspA. Therefore, in force-opposing experiments the observable step is that in which DNA moves relative to domain 2A while in force-assisting experiments the observable step is that in which DNA moves relative to domain 1A.

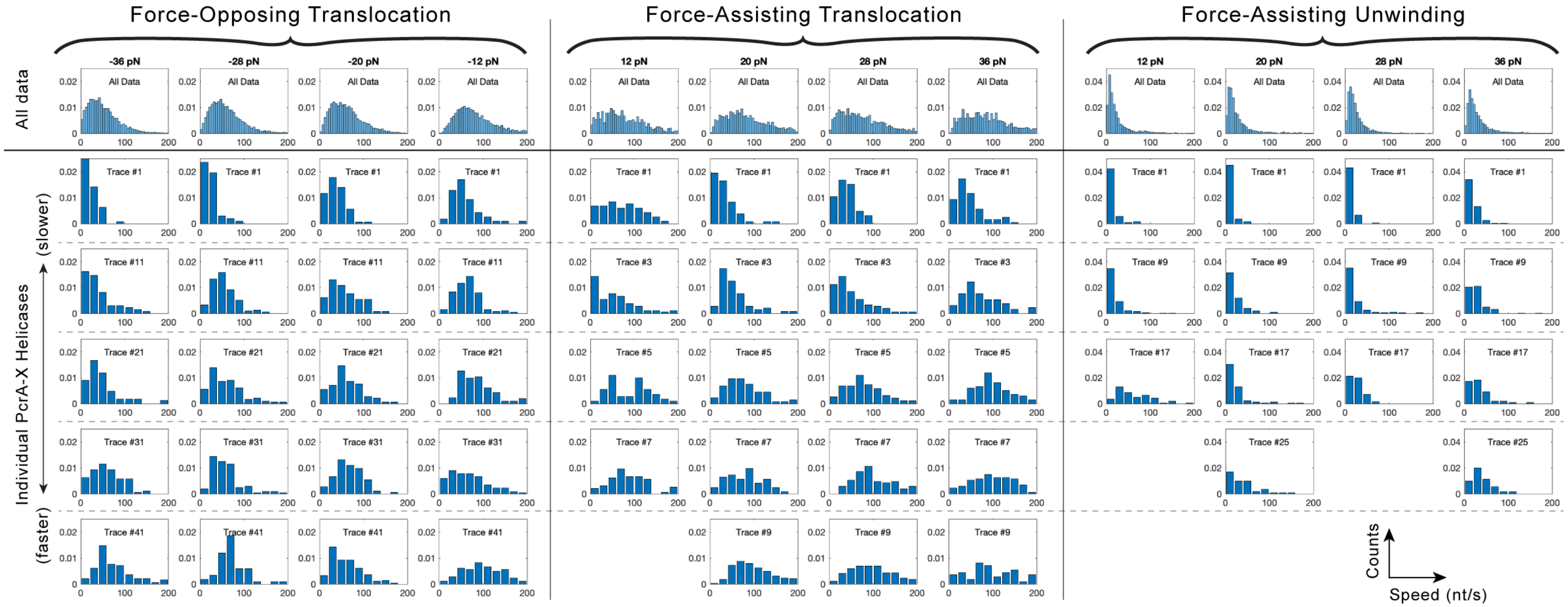

**Figure S11 | Static disorder among individual helicase reads.** Individual helicases had stepping rate histograms that differed significantly from the average histogram for all measurements. Below, several representative stepping rate histograms at saturating [ATP] in force-opposing translocation, force-assisting translocation, and force-assisted unwinding. The top row shows the histogram of all measured local stepping rates at that applied force for all helicase molecules, while lower rows show the range of stepping rates observed for an individual helicase as it traveled over several thousand bases as applied force was sequentially changed (as in Figure 1). As in the main text, local stepping rate measurements are made as sequential measurements of helicase stepping rate over 15 nt intervals along the DNA strand. Histograms are normalized as a probability distribution function so that the total bar area sums to 1 allowing direct comparison of the histograms with differing bin-widths. Here individual helicases have been sorted from slowest (top) to fastest (bottom) showing the range of stepping rates exhibited. Generally, individual helicase molecules are uniformly faster or slower than the average across all applied forces. Trace #'s relate only to helicase molecules within that dataset, i.e. Trace #1 in each configuration represents three separate helicases.

Michaelis-Menten fits and ADP-inhibition curves.

A

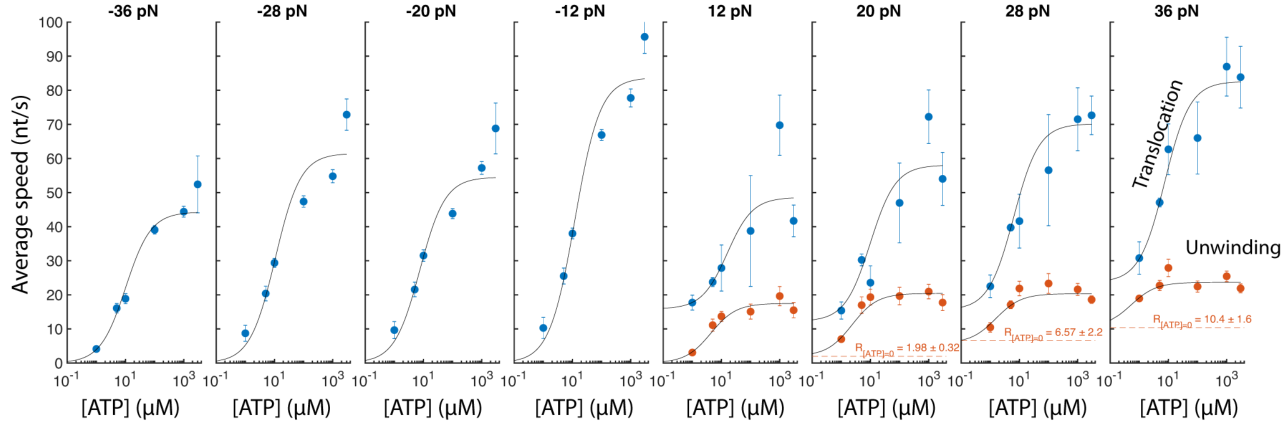

| | | | | | | | | |
|---|---|---|---|---|---|---|---|---|
| $K_M$ (μM) | 10.1 ± 1.6 | 10.7 ± 2.8 | 7.5 ± 2.1 | 12.0 ± 2.6 | 16 ± 12 | 9.8 ± 8.8 | 6.4 ± 1.8 | 7.5 ± 2.8 |
| Vmax (nt/s) | 44.3 ± 3.6 | 61.5 ± 5.9 | 54.5 ± 5.1 | 83.7 ± 6.1 | 32.7 ± 9.0 | 46 ± 15 | 54.7 ± 6.2 | 59.0 ± 8.2 |
| C (nt/s) | - | - | - | - | 15.8 ± 4.1 | 12.2 ± 7.5 | 15.5 ± 4.8 | 23.6 ± 6.6 |

Translocation

$$V = \frac{V_{max} \bullet [ATP]}{K_m + [ATP]} + C$$

| | | | | |
|---|---|---|---|---|
| $K_M$ (μM) | 4.33 ± 0.80 | 2.33 ± 0.61 | 1.5 ± 1.1 | 0.52 ± 0.32 |
| Vmax (nt/s) | 17.5 ± 1.5 | 18.5 ± 1.4 | 14.6 ± 3.4 | 13.3 ± 2.7 |
| C (nt/s) | 0* | 1.95 ± 0.34 | 5.7 ± 3.2 | 10.3 ± 2.4 |

Unwinding

B

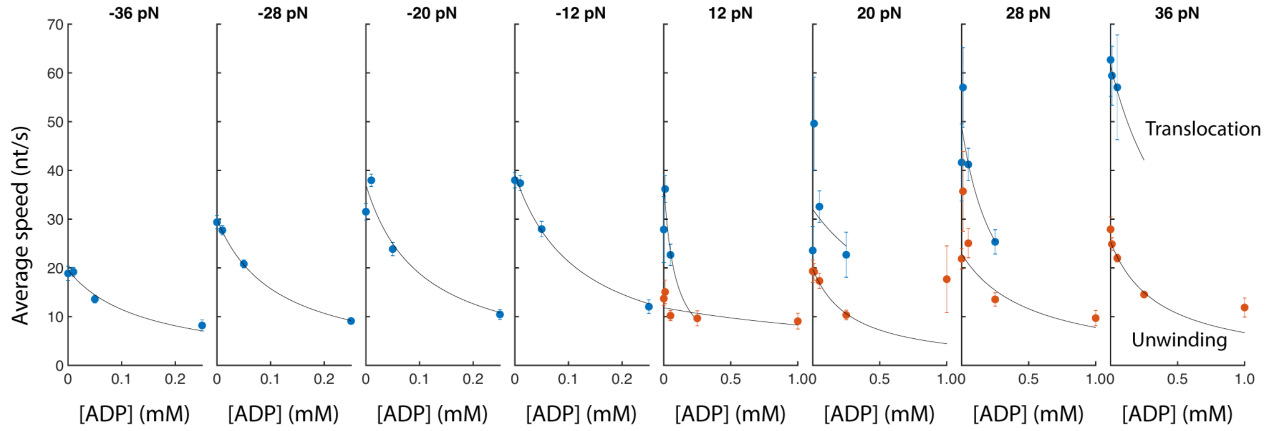

| | | | | | | | | |
|---|---|---|---|---|---|---|---|---|
| m (ms/μM) | 0.36 ± 0.12 | 0.301 ± 0.012 | 0.26 ± 0.13 | 0.215 ± 0.031 | 0.33 ± 0.33 | 0.039 ± 0.086 | 0.076 ± 0.025 | 0.030 ± 0.032 |
| b (ms) | 51.2 ± 3.9 | 33.39 ± 0.50 | 27.0 ± 3.7 | 25.51 ± 0.87 | 26.5 ± 8.2 | 31.2 ± 7.8 | 20.4 ± 2.5 | 16.25 ± 0.61 |

Translocation

$$V = \frac{1}{m \bullet [ADP] + b}$$

| | | | | |
|---|---|---|---|---|
| m (ms/μM) | 0.036 ± 0.040 | 0.175 ± 0.051 | 0.085 ± 0.038 | 0.109 ± 0.027 |
| b (ms) | 84.5 ± 9.6 | 50.5 ± 4.5 | 43.7 ± 6.5 | 39.5 ± 2.8 |

Unwinding

**Figure S12 | Michaelis Menten fits and ADP inhibition curves.** A) Michaelis-Menten fits for all applied forces for both translocation and unwinding experiments. Because motion was observed in the absence of ATP during force-assisted unwinding, a modified Michaelis-Menten equation is used to fit the data. C represents the stepping rate of PcrA driven by the assisting force rather than ATP hydrolysis. In this form, $V_{max}$ represents the maximum rate of ATP-driven motion not the asymptotic rate at saturating [ATP]. The asymptotic rate at saturating [ATP] is given by the sum of $V_{max}$ and C and the ratio of C to $V_{max}$ represents the ratio of forwards steps that are force-driven to those that are ATP driven. At 12 pN, the fit for C was fixed at the lower bound 0 suggesting force-driven unwinding occurs rarely at low assisting force. In unwinding experiments, as assisting force increases, only the [ATP]-independent stepping rate increases with force while $V_{max}$ and $K_M$ remain mostly unchanged, suggesting that the applied assisting force does not affect the rate-limiting step for on-pathway motion.

The rate increase observed for assisting force is then simply due to an increase in the off-pathway diffusive motion of the helicase driven by the assisting force. In the three rightmost graphs, the orange dashed line represents the observed stepping rate for unwinding at [ATP] = 0. B) ADP inhibition curves fit to a linear inhibition model. The data is consistent with competitive inhibition in which ATP and ADP compete for the ATP binding site.

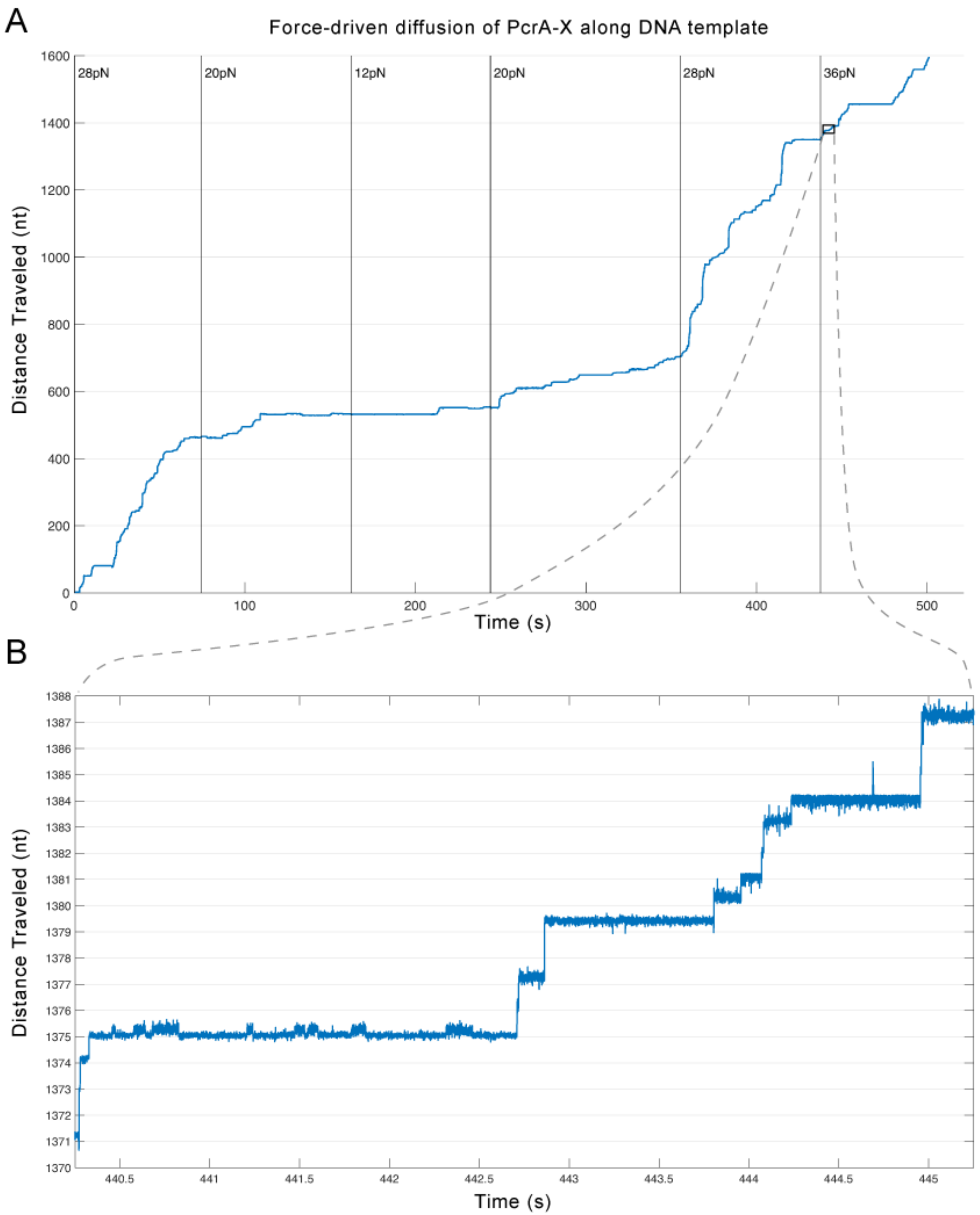

**Figure S13 | Stepping of PcrA helicase with no ATP present.** Under assisting forces without ATP, PcrA undergoes force driven unwinding. A) An example trace of PcrA undergoing force-driven unwinding with no ATP present. Various forces are applied ranging from 12pN to 36pN. This force-driven unwinding occurs at forces above 12 pN. Unwinding occurs at a low rate at low forces and is faster at higher forces. B) Zoom in detail of A) showing several single-nucleotide steps. Steps larger than 1nt are apparent in the data, we believe this [ATP]-independent motion reflects force-driven 1D diffusion of PcrA along the DNA backbone 'lattice.'

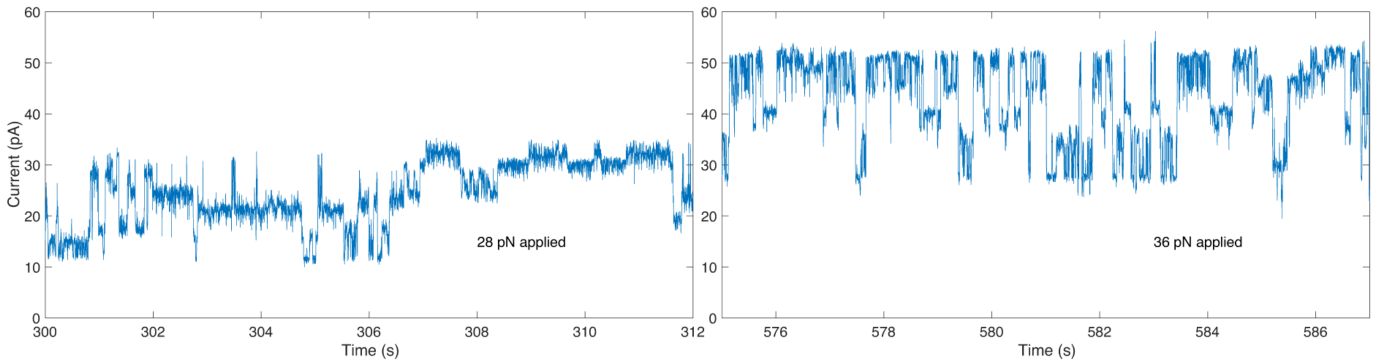

**Figure S14 | Backstepping at high [ADP] at 28pN and 36pN opposing force.** Representative ion current traces showing the high prevalence of backstepping at 28pN (left) and 36pN (right). Higher opposing forces with [ATP]=10μM and [ADP]=1000μM resulted in significant backstepping to the extent that automated alignment of ion-current levels to the ion-current consensuses (methods, Figs. S4-S6) was impossible.

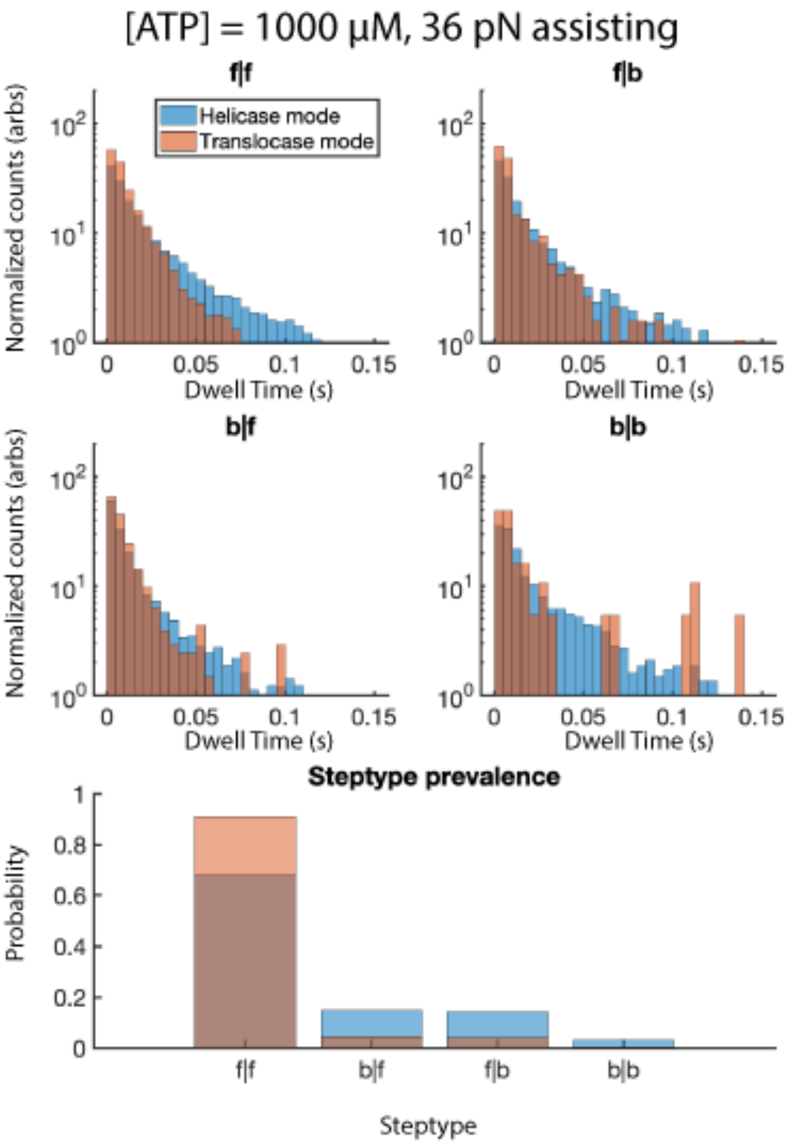

**Figure S15 | Dwell time histograms and prevalence of observed step types** at [ATP] = 1000 µM with 36 pN assisting force. Because step dwell times depend on the initial and terminal state of the enzyme system, analysis of SPRNT data must be separated into conditional step types: f|f (forwards steps that follow forwards steps), b|f (backwards steps that follow forwards steps), f|b (forwards steps that follow backwards steps), and b|b (backwards steps that follow backwards steps). Dwell time histograms for both helicase and translocase mode data are shown at 36 pN assisting force for each of the possible conditional step types. As is shown in the main text, f|f steps during unwinding are significantly longer than those observed during ssDNA translocation. Interestingly, the dwell time histograms for f|b steps for both unwinding and translocation are nearly identical suggesting the presence of a DNA duplex does not affect this portion of the mechanochemical pathway. This is consistent with the model shown in Figure 2 of the main text wherein duplex unwinding occurs after ATP hydrolysis. Backstepping observed here occurs before ATP hydrolysis and is a simple backtracking of domain 1A while ATP remains bound. Presence or absence of a DNA duplex should have no effect on this kinetic step.

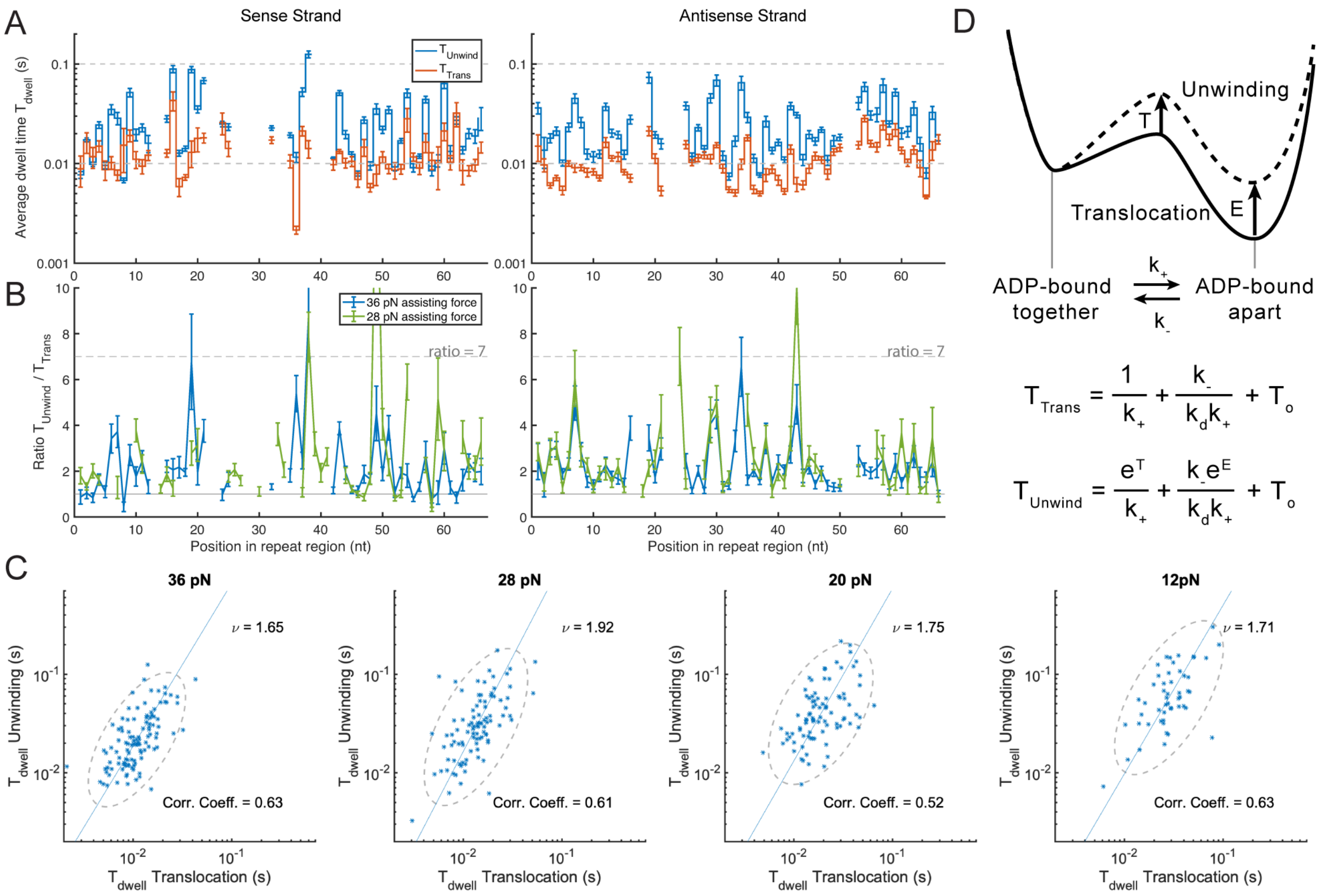

**Figure S16 | Sequence dependent translocation and unwinding for repeat DNA.** A) Average dwell-time as a function of location within the repeat sequence for the sense strand (left) and antisense strand (right). Unwinding dwell-times in blue and translocation dwell-times in orange. B) The ratio of $T_{Unwind}/T_{Trans}$ for both sense and antisense strands. In the Betterton model[11], this ratio for a passive helicase would be 7, for an active helicase the ratio would be 1. Depending on the sequence context, PcrA displays a range of "activity." C) $T_{Trans}$ vs. $T_{Unwind}$ at 36 pN, 28 pN, 20 pN and 12 pN assisting force for the dwell times shown in Figure 4A of the main text plotted on a log-log scale. Dwell-time pairs for both sense and antisense sequences have been combined into one plot. The correlation coefficient between $\log(T_{Trans})$ and $\log(T_{Unwind})$ is 0.63, 0.61, 0.52, and 0.63 respectively. The correlation between $T_{Trans}$ and $T_{Unwind}$ suggests that duplex unwinding is simultaneous with the advance of domain 2A in our model (Fig. 3) rather than introducing a new rate-limiting step. D) A model using transition state theory for how the presence of the duplex modifies PcrA stepping kinetics. Both the transition state and the energy of the second bound state are increased by T and E (units of kT), respectively, in response to the presence of a duplex. At saturating [ATP] and with [ADP]=0 the equation for the average dwell-time of a step in the model depicted in Figure 2 can be solved for analytically[9]. $k_-$ and $k_+$ are the backwards and forwards rates of domain 2A for PcrA translocation, $k_d$ is the rate of ADP dissociation, and $T_o$ is the combination of all other rates unaffected by the presence of the duplex (ATP binding, hydrolysis, and motion of domain 1A). In our model, $k_-$, $k_+$, T, and E are all sequence dependent. $k_-$ and $k_+$ are determined by the ssDNA sequence whereas T and E are dependent upon the duplex sequence. The leftmost term represents how the forwards rate, domain 2A advance, is modified by the duplex whereas the middle term represents how the equilibrium between the two states on either side of the kinetic step is modified.

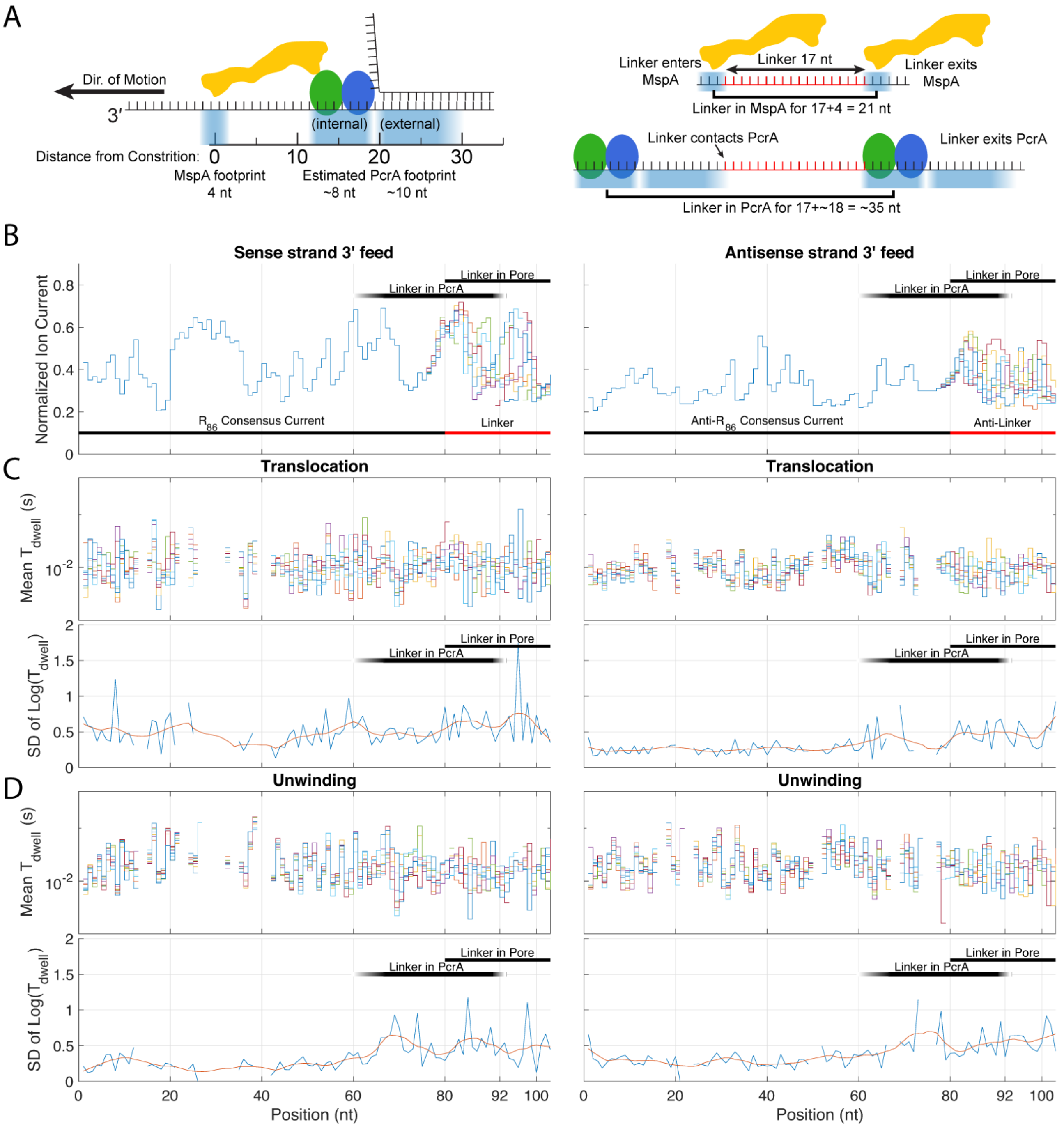

**Figure S17 | Determination of where sequence-dependent effects are localized.** A) Schematic showing PcrA at rest on top of MspA during an unwinding experiment. Ion currents in MspA are affected by 4 bases in and around the constriction. As such, the linker footprint within the pore will be ~20 nucleotides in which linker DNA affects the ion-currents observed. PcrA is expected to be in contact with nucleotides ~12-30 upstream from the MspA constriction, where "internal" contacts are those within the ssDNA channel bound to RecA-like walker domains and "external" contacts are those within the dsDNA duplex that contact the exterior of PcrA in the crystal structure 3PJR. Thus, PcrA will be in contact with some portion of the 17-base linker region for ~18 + 17 = ~35 steps. B) Consensus current for sense and antisense strand demonstrating the effect of the linker region when it is within the pore constriction. C) *top:* Average translocation dwell-times corresponding to the current levels shown above in B). *Bottom:* Standard deviation of the log dwell times shown above (blue). Smoothed with a 10-nt box filter (red). This is a proxy for the spread in observed dwell times for each instance of each repeat. An increase in this value indicates that the dwell-times for this level are inconsistent. D)

Same as C) for Unwinding. The physical separation of PcrA and the pore constriction allows for localization of sequence dependent effects. There is a marked increase in SD(log($T_{dwell}$)) approximately ~15 nucleotides prior to the linker regions' entry into the MspA constriction. Put simply, when the same sequence is in MspA but different sequence is in in PcrA, dwell-times are significantly different. This demonstrates unequivocally that different DNA bases within PcrA are responsible for sequence-dependent effects. Unfortunately proving the converse (that bases within the pore constriction alone do not affect step kinetics) with the above data is complicated by the difficulty of aligning current levels in the linker regions to one another. Small misalignments of a few levels within each linker consensus can also give rise to large variation in $T_{dwell}$. However, previous SPRNT experiments with Hel308 found DNA within the pore constriction was not responsible for sequence-dependent kinetics.

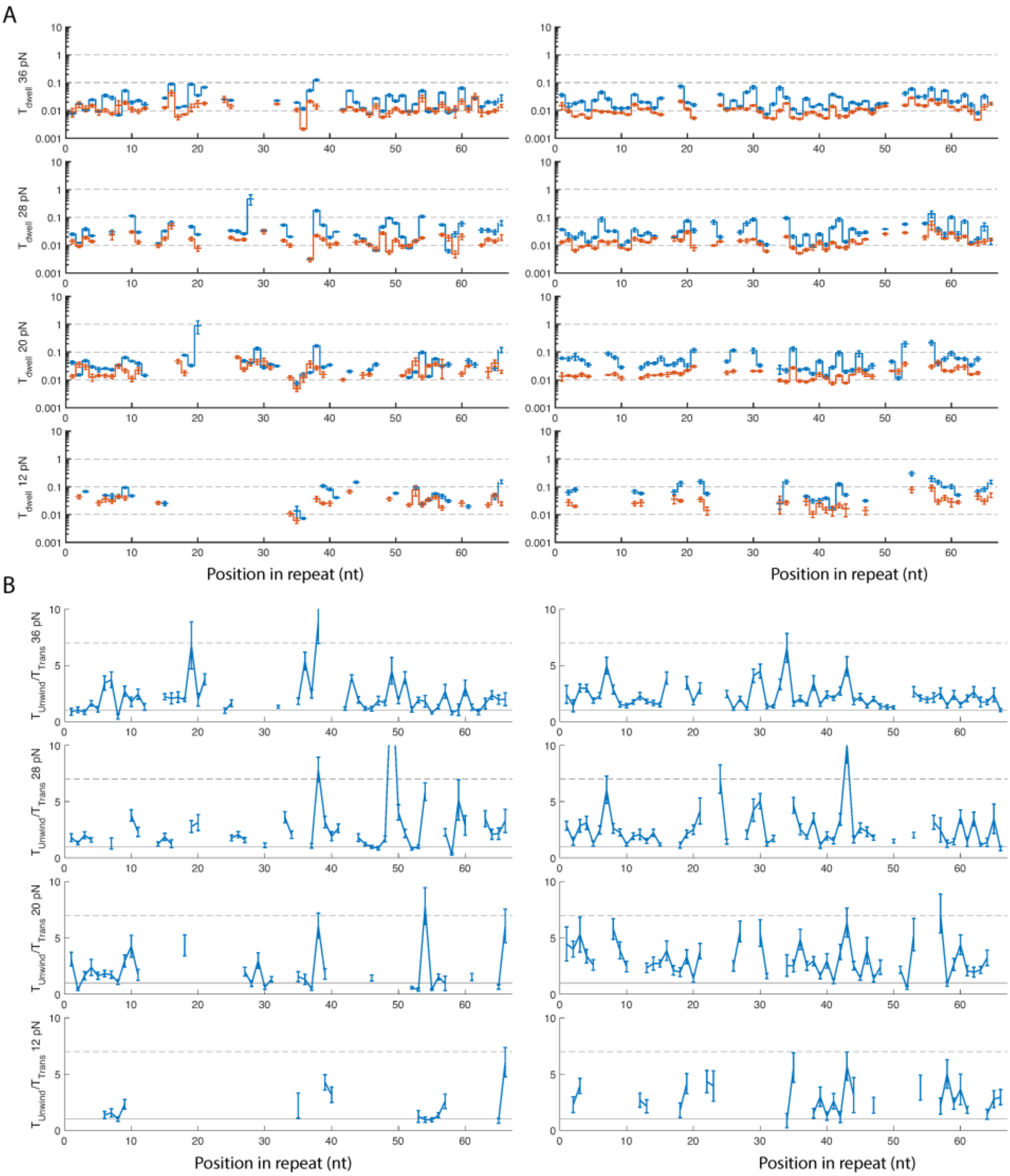

**Figure S18 | Sequence-dependent kinetics at all applied forces.** A) The median dwell time for each location along the repetitive DNA section for translocation (orange) and unwinding (blue) at various assisting forces. Sense strand data is at left while antisense strand data is at right. We exclude data in which the linker region is within the pore or PcrA. There is over an order of magnitude variation in median dwell time depending on the particular sequence context within the PcrA helicase. This means that only a few particularly slow sequence contexts end up dominating the average behavior of the helicase. B) The ratio of average unwinding dwell times to translocation dwell times from A). The ratio reveals a sequence-dependent pattern in both sense and antisense sequences across all applied forces. This ratio indicates how much slower unwinding is in comparison to translocation for each location along the DNA sequence and reveals the component of the sequence-dependent signal in A) that is due to the presence of the DNA duplex. The applied force does not appear to affect the ratio significantly which is consistent with our model of how force affects PcrA motion in SPRNT. Put another way, the applied force does not appear to affect the duplex unwinding step.

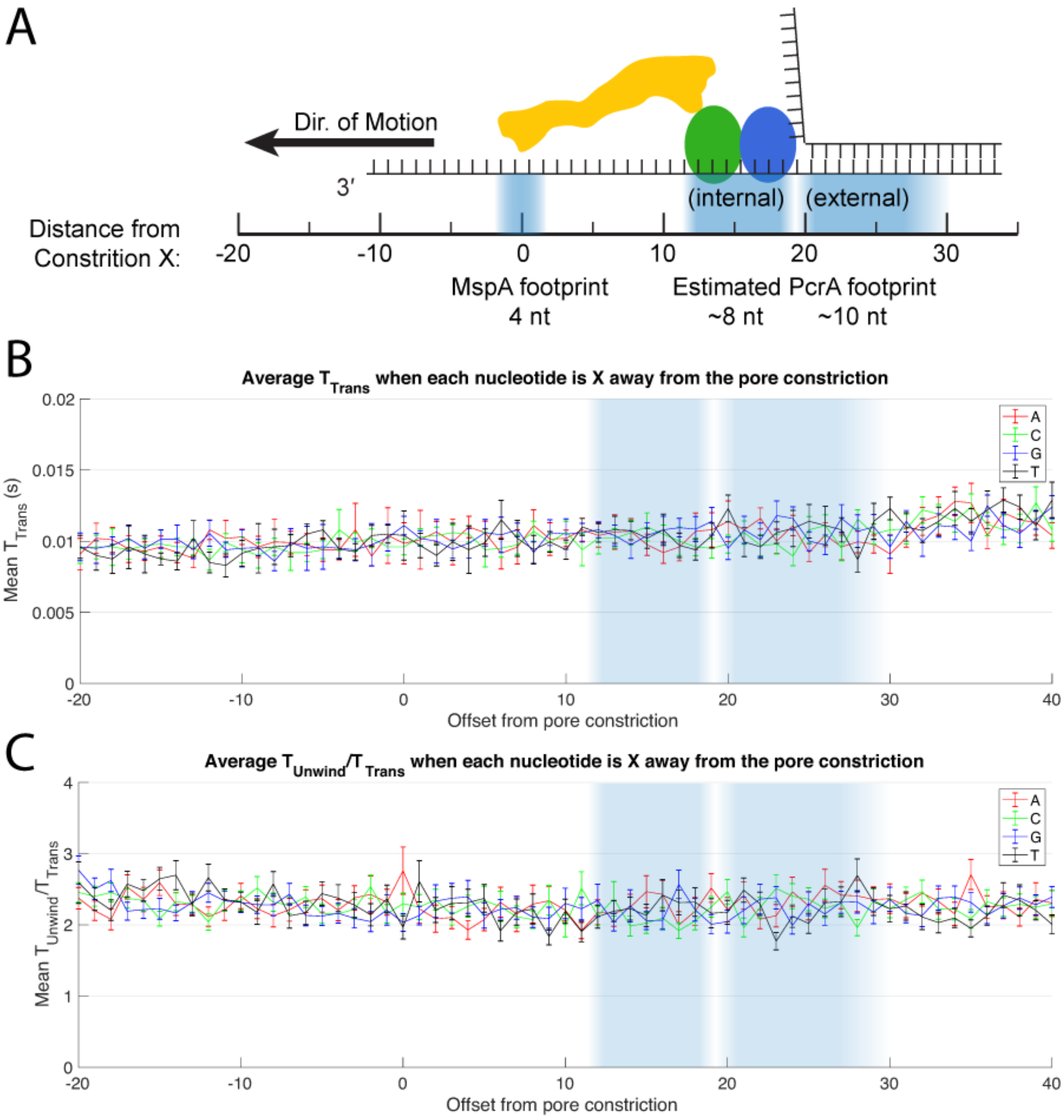

**Figure S19 | Analysis of the effect of nucleotides on $T_{dwell}$.** A) Schematic of PcrA sitting on top of MspA during an unwinding experiment demonstrating the x-axis used in the following graphs. B) Average dwell-time when A, C, G, or T (red, green, blue, and black, respectively) are x nucleotides from the pore constriction. For example, the black datapoint at X=30 represents that when T is located 30 nucleotides upstream of the pore constriction the average dwell-time is ~12.5 ms. PcrA is expected to reside ~12 to 30 nucleotides upstream of the MspA constriction. C) The average ratio $T_{Unwind}/T_{Trans}$ when A, C, G, or T (red, green, blue, and black, respectively) are X-nucleotides away from the MspA constriction. Graphs in both B) and C) show little average effect of a single nucleotide on the average dwell time or ratio $T_{Unwind}/T_{Trans}$. This indicates that more than just a single site is responsible for the sequence-dependent kinetics observed in this study.

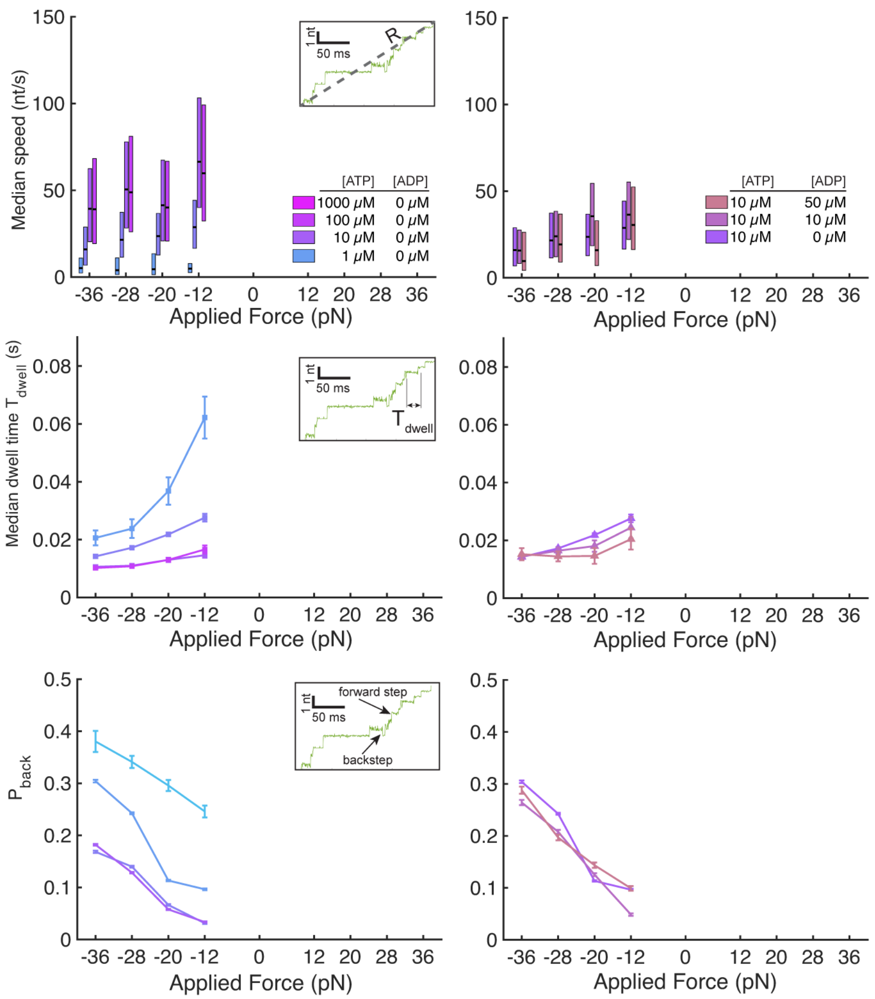

**Figure S20 | Average kinetic behavior for uncrosslinked PcrA (C96A, C247A, N187C and L409C)**. just as in Figure 3 of the main text. Data shown is for force-opposing translocation. Uncrosslinked PcrA showed little unwinding activity thus there was insufficient data to compare uncrosslinked PcrA to PcrA in force-assisted configurations. The data above shows similar force, [ATP], and [ADP] dependence to PcrA data shown in the main text suggesting that the act of crosslinking PcrA is not responsible for the data described in the main text.

**PcrA overexpression, purification, and crosslinking protocol.**

PcrA expressing vector: This protocol describes the purification and crosslinking of *Bacillus stearothermophilus* PcrA helicase. Our PcrA mutant has N terminal tag containing 6xHis-tag. Our PcrA mutant has two native cysteines C96A and C247A removed, while N187C and L409C are introduced for crosslinking. Briefly, we inserted PcrA sequence between NdeI and BamHI sites of pET-11b vector and transformed it into E. coli BL21(DE3) cells and grown in TB (Terrific broth) medium. TB increases the cell mass and has no negative effect on the quality and overexpression levels of our protein.

Purification columns: Our purification protocol uses a gravity flow column. The first purification stage involves Ni-NTA column, while the second stage is the single-stranded DNA cellulose column that can be also combined with the crosslinking step.

Bacterial expression media: We use granulated Terrific Broth from Invitrogen containing 11.8 g SELECT Peptone 140, 23.6 g Yeast Extract, 9.4 g dipotassium hydrogen phosphate, 2.2 g potassium dihydrogen phosphate per 1 liter. Catalog number 22711022.

Buffers and other chemicals:

Lysis Buffer: (50 mM Tris, 5 mM Imidazole, 200 mM NaCl, 20% (w/v) Sucrose, 15% (v/v) Glycerol, Adjust to pH 7.6 with HCl at RT; filter (0.22 µm); prepare 0.5 l.

Buffer A: 50 mM Tris, 5 mM Imidazole, 150 mM NaCl, 25% (v/v) Glycerol, adjust to pH 7.6 with HCl at RT, filter (0.2 µm); prepare 2 l.

Buffer A1M: 50 mM Tris, 5 mM Imidazole, 1 M NaCl, 25% (v/v) Glycerol, Adjust to pH 7.6 with HCl at RT, filter (0.2 µm); prepare 0.5 l.

Buffer B: 50 mM Tris, 1 mM EDTA, 20% (v/v) Glycerol, adjust to pH 7.6 with HCl at RT, filter (0.2 µm); prepare 2 l

Buffer B2M: 50 mM Tris, 1 mM EDTA, 2M NaCl, 20% (v/v) Glycerol, adjust to pH 7.6 with HCl at RT, filter (0.2 µm); prepare 2 l

Imidazole Elution Buffer:200mM imidazole in Buffer A, pH 7.4 using HCl or NaOH; filter (0.2 µm); prepare 0.5 l.

Buffer B100mM: Mix Buffer B and B2M in the ratio 19x buffer B + 1X Buffer B2M

B1M is ssDNA-cellulose elution buffer: by mixing Buffer B and B2M in the ratio 1 to 1

PMSF (phenylmethylsulfonyl fluoride 17.5mg/ml in isopropanol)

Lysozyme (20mg/ml in water)

TCEP, tris(2-carboxyethyl)phosphine Use 0.5M, pH 7.0 from ThermoFisher

BM(PEG)2 (1,8-bismaleimido-diethyleneglycol) crosslinker 10 mM in DMF (dimethylformamide)

Ampicillin stock: 100 mg/mL in deionized water

Chloramphenicol stock: 30 mg/mL in ethanol

Protein overexpression: We transform vector into E. coli BL21 (DE3) and plate on LB agar with 100 µg/mL Ampicillin and 30 µg/mL Chloramphenicol at 37°C overnight. We then pick a single colony and grow it in LB medium containing 100 µg/mL Ampicillin and 30 µg/mL Chloramphenicol at 37°C until $OD_{600}$ reaches 0.5. A large 4L flask containing 1L of TB medium with 100 µg/mL Ampicillin and 30 µg/mL Chloramphenicol is inoculated with the small culture and grown at 37°C. When $OD_{600}$ reaches about 0.3, the temperature is lowered to 18°C and grown until $OD_{600}$ is 0.8. We then induce overexpression by adding 0.5 mM IPTG and grow overnight. We harvest cells the next day by centrifugation in Beckman/Coulter JA-14 rotor, 15min, 5000 rpm at 4°C. The pellet may also be stored at -80 °C for future purifications.

Cell lysis: We first measure the mass of the cell pellet and per 1 gram of pellet we add 100 $\mu$L of lysizymes and 6 $\mu$L of PMSF. For 5 grams of pellet we add 40 mL of lysis buffer and sonicate using Fisher Sonic Dismembrator 500 . The total time is set to 5 minutes with 20% of max power and 0.5s ON and 0.5 OFF duty cycle. Once the pellet is fully resuspended and has a milk-like consistency we stop he sonication. One usually does not have to wait 5 minutes to complete the sonification. Protein expression levels can be checked with the SDS-PAGE analysis.

Centrifugation: Resuspended and lysed cells require centrifugation at 14,000 RPM and 4°C for at least 30 minutes. (1 hour is fine too) Pellet is discarded and supernatant containing soluble protein is transferred to a new 50 mL tube. Supernatant containing PcrA is now ready to be mixed with Ni-NTA column for the affinity-based purification.

Ni-NTA column equilibration: Purchase Ni-NTA agarose resin contains the storage buffer and must be equilibrated with Buffer A + 5 mM TCEP prior to adding it to the supernatant containing PcrA. We use 1 mL of Ni-NTA resin per 1 gram of cell pellet. Ni-NTA agarose resin is well mixed with 40 mL of buffer A + 5 mM TCEP in 50 mL tube, then centrifuged down at 1500 RPM for 10 s. The supernatant is removed, and resin washed by repeating the procedure 2 more times. Altogether 3 washes with 40 mL of buffer.

Batch mixing with Ni-NTA agarose resin: Washed Ni-NTA agarose resin is mixed with ~40 mL of cell supernatant and stir mixed at 4°C for 90 minutes. Alternatively, one can mix at room temperature for 30 minutes, because PcrA is stable at RT. During this step PcrA from the solution binds to Ni-NTA beads. Ni-NTA column has a light blue color, but after PcrA binding, beads change color towards white/yellow. One should avoid aggressive shaking of protein-bound Ni-NTA column. Gentle inverting of the tube is preferred.

Washing protein-bound resin: After stir mixing for 90 minutes, we spin down the tube at 1500 RPM for 10 s and discard the supernatant. After the centrifugation, one should make sure that the beads are at the bottom of the 50 ml tube. After removing the supernatant, we add additional 40 mL of Buffer A + 5 mM TCEP, gently invert the tube several times until the bead-resin is fully resuspended and spin down again. The batch washing procedure is repeated 3 times.

Gravity column wash: Protein-bound resin is transferred to a 20-ml disposable chromatography column (BioRad). Since the resin is hard to pour after the centrifugation, we resuspend it in 10 mL of Buffer A + 5 mM TCEP and then add to the column and we transfer as much beads as possible. The volume of the resin in column (CV) is typically 2 to 3 mL. We use CV as a volume unit for further wash steps. We add 15 CVs of Buffer A+5mM TCEP at 4°C and wait for it to flow through. Alternatively, we use a vacuum manifold attached to the chromatography column to speed up the wash. The next washing step is with 10 CV of A1M buffer containing 1M NaCl. Washing protein with the high salt releases any DNA or RNA bound to the protein. Finally, we wash PcrA with 20 CV of Buffer A without TCEP.

Protein labeling (optional): If there is a need to label the protein with maleimide-fluorophore, labelling can be carried out while the protein is bound to the column. Excess dye can be washed with Buffer A.

Elution: We add 2 CV of imidazole elution buffer to the chromatography column and collect several 0.5 mL fractions. As fractions are collected, we periodically add more elution buffer to the column and measure the eluted protein concentration on Nanodrop. In case the concentration exceeds 4 mg/ml, we add Buffer A to dilute the eluted protein below 4 mg/ml (50 $\mu$M) and prevent aggregation. Elution fractions can be checked on SDS-PAGE gel for purity. Fractions can be stored at 4 degrees for a few days before the next step is carried out. At this stage, the activity of the protein can be tested at 37 degrees, using a FRET-pair labeled 18-mer with 18-poly-dT on 3' end.

Single-stranded (ss) DNA-cellulose purification and crosslinking: The imidazole elution provides a mixture of PcrA protein, including properly folded, misfolded, active, and inactive protein. We want to select only active protein that can bind the single-stranded DNA and for this purpose we use ssDNA-cellulose column. We use Deoxyribonucleic acid−cellulose single-stranded from calf thymus DNA purchased from Sigma-Aldrich (Item D8273-5G).

Hydrating ssDNA cellulose column: Single-stranded DNA purification step is carried out in a 20-ml chromatography column at 4 degrees. We first close the bottom of the column to prevent the flow and fill the column with 15 mL of buffer B2M. We then sprinkle in ssDNA-cellulose powder and let it settle at the bottom of the column. When the volume of sprinkled ssDDNA-cellulose is about 2 ml, we stop sprinkling and open the bottom to let the buffer flow. We let buffer B2M flow through for about 1 hour to hydrate and wash the column. Buffer flowing through this column is slow, so the process should be started early in the day. After 1 hour of hydration, we wash the column with 20 CV or B100mM (low salt buffer).

Loading eluted fractions. When the column is ready, eluted fractions are loaded to it. Only active protein will bind, while the rest will flow through. Save the flow through for SDS-PAGE analysis. It is also recommended to check the concentration on the nanodrop and detect when the column is saturated by PcrA. (When it is saturated, 100% of added protein flows through)

Option A for PcrA without crosslinking: We wash the PcrA-loaded column with 10 CV of buffer B100mM to remove the excess imidazole. Once wash buffer has flown through, we elute with buffer B1M. Under the high-salt buffer B1M condition, PcrA dissociates from the ssDNA column. We collect 0.5 mL elution fractions and check the concentration. After this step, no dialysis is required, unless there is a need to concentrate the sample. We add glycerol and TRIS to make the final concentration of the storage buffer: 50% glycerol, 50 mM TRIS pH 7.6, 600 mM NaCl. We split PcrA in small tubes, measure concentration, and store at -80 or -20 degrees.

Option B for crosslinked PcrA: We wash the column with 10 CV of B100mM + 5mM TCEP. This step eliminates excess imidazole from the solution and TCEP exposes cysteines for crosslinking, followed by another wash with 10 CV of B100mM (no TCEP). We dissolve BM(PEG)2 (1,8-bismaleimido-diethyleneglycol) to 10 mM in DMF, and then further dilute to 100 $\mu$m in B100mM. The column is washed with B100mM + BM(PEG)2 at room temperature for 1 hour. We then wash with 10 CV of B100mM to remove the excess crosslinker and elute PcrA with B2M. Finally, we check the concentration on nanodrop and add glycerol and TRIS to make the final concentration of the storage buffer: 50% glycerol, 50 mM TRIS pH 7.6, 600 mM NaCl. We split PcrA in small tubes and store at -80 or -20 degrees. This method has higher crosslinking efficiency, see gel lanes #5 and #6.

Alternative crosslinking: Alternatively, crosslinking can be carried out after eluting in Option A by adding crosslinker to a free protein in solution and incubation for 2 hour at room temperature. Removing excess crosslinker in that case require dialysis against the storage buffer: 50% glycerol, 50 mM TRIS pH 7.6, 600 mM NaCl. The crosslinking efficiency of this method is lower, see gel lanes #2 and #3.

| | |
|---|---|
| | Lanes:<br>1. Protein Ladder<br>2. Low efficiency crosslinking<br>3. Low efficiency crosslinking<br>4. Left empty<br>5. High efficiency crosslinking<br>6. High efficiency crosslinking |

**Figure S21 | SDS-PAGE gel of crosslinked PcrA.** Loading PcrA-X on the gel results in 3 bands. Uncrosslinked PcrA-X is linear after denaturing; hence it has the highest mobility and contributes to the bottom-most band. Crosslinked PcrA-X contributes to the middle band. Crosslinking between two or more separate PcrA molecules creates a multimer and contributes to the top-most band. Efficient crosslinking has the highest intensity of the middle band. Lane 1 is the protein ladder. Lanes 2 and 3 shows low crosslinking efficiency. Lane 4 is left empty. Lanes 5 and 6 show high crosslinking efficiency obtained with Option B.

**Additional Sources Cited.**
